# Hierarchical Heterogeneity Across Human Cortex Shapes Large-Scale Neural Dynamics

**DOI:** 10.1101/341966

**Authors:** Murat Demirtaş, Joshua B. Burt, Markus Helmer, Jie Lisa Ji, Brendan D. Adkinson, Matthew F. Glasser, David C. Van Essen, Stamatios N. Sotiropoulos, Alan Anticevic, John D. Murray

**Author notes:** **Corresponding Author:** John D. Murray, Department of Psychiatry,Yale School of Medicine, 40 Temple Street, Suite 6E, New Haven, Connecticut, 06510, USA.

## Abstract

The large-scale organization of dynamical neural activity across cortex emerges through long-range interactions among local circuits. We hypothesized that large-scale dynamics are also shaped by heterogeneity of intrinsic local properties across cortical areas. One key axis along which microcircuit properties are specialized relates to hierarchical levels of cortical organization. We developed a large-scale dynamical circuit model of human cortex that incorporates heterogeneity of local synaptic strengths, following a hierarchical axis inferred from MRI-derived T1w/T2w mapping, and fit the model using multimodal neuroimaging data. We found that incorporating hierarchical heterogeneity substantially improves the model fit to fMRI-measured resting-state functional connectivity and captures sensory-association organization of multiple fMRI features. The model predicts hierarchically organized high-frequency spectral power, which we tested with resting-state magnetoencephalography. These findings suggest circuit-level mechanisms linking spatiotemporal levels of analysis and highlight the importance of local properties and their hierarchical specialization on the large-scale organization of human cortical dynamics.

## Introduction

The spatiotemporal dynamics of a neural system are shaped by structural constraints on interactions among the system’s components, as well as the intrinsic dynamical properties of those components. An important open question in systems neuroscience is how areal heterogeneity of local circuit properties across cortex shapes large-scale structure-function relationships. Hierarchical organization provides a parsimonious principle for the anatomical properties of long-range inter-areal connections in primate cortex (Felleman and Van Essen, 1991; Dombrowski et al., 2001; Markov et al., 2014). Anatomically-defined cortical hierarchy has been found to align with sensory processing hierarchies, with early sensory areas at lower levels and association areas at higher levels (Felleman and Van Essen, 1991; Markov et al., 2014). Furthermore, functional and dynamical properties (Murray et al., 2014; Honey et al., 2012), as well as with specialization of cortical microcircuitry (Burt et al., In press; Chaudhuri et al., 2015), vary across hierarchical levels. Yet it is unclear how hierarchical specialization of local circuit properties across human cortex shapes the large-scale organization of neural dynamics.

Advances in magnetic resonance imaging (MRI) have provided noninvasive methods for characterizing large-scale connectivity in the human brain at the structural and functional levels. Structural connectivity (SC) is often inferred from diffusion MRI (dMRI), which aims to quantify the density of anatomical fibers linking brain regions (Le Bihan and Iima, 2015). The functional organization of activity in the human brain has been studied most extensively through functional MRI (fMRI) measurements of blood-oxygen-level-dependent (BOLD) signals across the brain (Biswal et al., 2010). Resting-state functional connectivity (rs-FC) provides a measure of the temporal correlations in spontaneous activity between brain areas, and has revealed an intrinsic architecture of the human brain (Cole et al., 2014). There is a substantial literature establishing relationships between dMRI-based SC and fMRI-based rs-FC (e.g., Greicius et al., 2009; Hagmann et al., 2008). Recent findings suggest that hierarchical organization may be a useful principle for describing rs-FC patterns in human cortex (Margulies et al., 2016), including capturing sensory–association differences in inter-individual variation (Mueller et al., 2013; Finn et al., 2015) and dysfunction in disease states (Baker et al., 2014; Yang et al., 2016).

Computational models of large-scale brain circuits propose dynamical circuit mechanisms linking the structural and functional organization of human cortex. In a major class of biophysically-based dynamical models, large-scale patterns of rs-FC arise through physiological dynamics of local cortical circuits interconnected through long-range structural connections (Breakspear, 2017; Deco et al., 2011). Large-scale computational modeling studies have found that simulated rs-FC in a biophysically-based circuit model can capture empirical rs-FC patterns better than the SC alone. Importantly, simulated FC in these models is shaped by the neurophysiological properties of the local circuits, such as strengths of excitatory and inhibitory synaptic connections (Deco et al., 2013; 2014b; Yang et al., 2014). However, the role of inter-areal heterogeneity of local circuit properties has not been systemically studied in large-scale models of human cortex.

Microcircuit specialization across human cortex can be informed by structural neuroimaging measures of cortical architectural variation. For instance, the MRI-derived contrast ratio of T1- to T2-weighted (T1w/T2w) maps has been proposed to provide an *in vivo* measure of intracortical myelin content (Glasser and Van Essen, 2011; Glasser et al., 2014). Cortical myelin content, measured by quantitative T1 mapping, has been observed to correlate with a prominent sensory–association gradient in rs-FC variation (Margulies et al., 2016; Huntenburg et al., 2017). Burt et al. (In press) found that the T1w/T2w map provides a noninvasive neuroimaging proxy measure of anatomical hierarchy in primate cortex. Furthermore, multiple aspects of hierarchical specialization, including in excitatory and inhibitory microcircuitry, vary along this cortical axis. In human cortex, the T1w/T2w map captures the dominant areal pattern of variation in gene expression (Burt et al., In press). We hypothesized that hierarchical specialization of local microcircuitry across human cortex, as captured by T1w/T2w maps, shapes the large-scale organization of rs-FC.

To address these issues, we developed a large-scale cortical circuit model incorporating hierarchical heterogeneity of local microcircuit properties, and quantitatively fit the model using multimodal human neuroimaging data from the Human Connectome Project (HCP) (Van Essen et al., 2013). We used the T1w/T2w map to parametrize hierarchical heterogeneity in local synaptic strengths across cortical areas. Compared to a model with homogeneous microcircuit properties across areas, this heterogeneous model better captured empirical rs-FC patterns, with the T1w/T2w map providing a preferential axis for areal heterogeneity compared to random heterogeneity maps. Furthermore, the model predicts areal heterogeneity in the spectral features of high-frequency neural dynamics, which we found to be consistent with resting-state magnetoencephalography (MEG). Our study provides a computational framework to study how areal specialization of microcircuitry shapes large-scale network function of the human brain, opening applications to neuropsychiatric disorders and pharmacological effects.

## Results

We first describe the computational framework for the large-scale circuit model of human cortex, incorporating areal heterogeneity of local properties, which we applied to the HCP multimodal neuroimaging dataset from a large number of healthy subjects (N=334) (**Figure 1A**). The cortical surface was parcellated into multiple contiguous areas. Here we applied a recently developed multimodal parcellation from the HCP which yielded 180 cortical areas per hemisphere (Glasser et al., 2016). Each cortical area was modeled as a local circuit comprising excitatory pyramidal neurons and inhibitory interneurons coupled through recurrent synaptic interactions, with neurophysiologically interpretable parameters governing local dynamics, as described below. Areas in the large-scale network interact via structured long-range excitatory projections constrained by a structural connectivity (SC) matrix, derived here from diffusion MRI (dMRI) and probabilistic tractography (**Figure S1)**. We simulated only within-hemisphere interactions, to focus model fitting on capturing the network structure of rs-FC (Deco et al., 2014a) (**Figure S1D**). The SC matrix thereby provides a structural scaffold for long-range neural interactions in the model.

**Figure 1:**
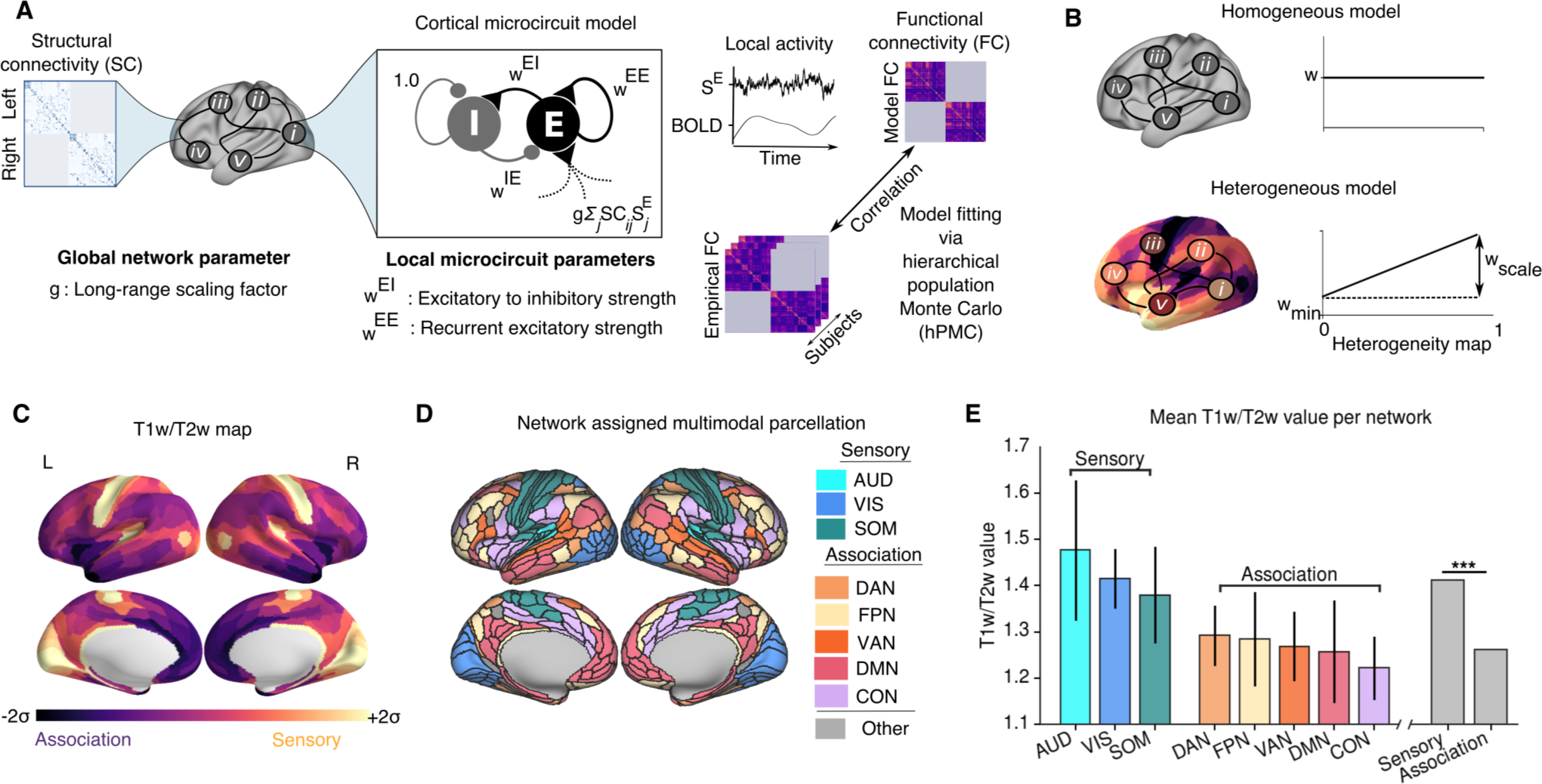
Large-scale Model of Human Cortex with Heterogeneous Local Circuit Properties. **(A)** Model framework. Each parcellated cortical area is modeled as coupled excitatory (E) and inhibitory (I) populations. Areas interact through long-range projections following dMRI-derived intra-hemispheric structural connectivity (SC). Fit model parameters comprised recurrent excitatory strength (*w^EE^*), excitatory-to-inhibitory strength (*w^EI^*), and a global coupling parameter scaling the strength of long-range connections (*g*). Inhibitory-to-excitatory strengths (*w^IE^*) were adjusted to to maintain a uniform baseline excitatory firing rate across areas. Dynamics of synaptic gating variable (*S^E^*) are transformed into a simulated BOLD signal via the Balloon-Windkessel hemodynamic model. For computational tractability of model fitting, model BOLD functional connectivity (FC) matrices were calculated via linearization of the extended dynamical equations around the fixed point of the system. Model parameters were fit to maximize the similarity between model and empirical FC matrices. **(B)** Parametrizing local properties via a heterogeneity map. In the homogeneous model, the parameters (*w^EI^* and *w^EE^*) were identical across cortical regions. In the heterogeneous model, the parameters (*w^EI^* and *w^EE^*) varied across cortical areas based on a heterogeneity map *h*, whose minimum and maximum value is 0 and 1, respectively. For each region (*i*) the parameter values were set by an affine function of the heterogeneity map values {*h_i_*}, characterized by an intercept *w_min_* and scale factor *w_scale_*: *w_i_* = *w_min_* + *w_scale_h_i_*. **(C)** Cortical T1w/T2w map. The median (*N* = 334 subjects) cortical T1w/T2w map values of each parcellated cortical area (180 per hemisphere). **(D)** Network assignments. Cortical areas were assigned to eight functional resting-state networks (RSNs) comprising three sensory (AUD, auditory; VIS, visual; SOM, somatomotor) and five association (DAN, dorsal attention; FPN, fronto-parietal; VAN, ventral attention; DMN, default mode; CON, cingulo-opercular) networks. **(E)** T1w/T2w map values per RSN, averaged across areas. T1w/T2w values are significantly lower in association RSNs than in sensory RSNs (*p* < 0.003, Wilcoxon signed-rank test, difference between sensory and association T1w/T2w across subjects). Error bars indicate the standard deviation across areas within an RSN.

The model simulates the time-varying activity of excitatory and inhibitory neuronal populations in a local circuit for each cortical area. For computational tractability of model fitting, as well as mathematical analysis of the system, we used a reduced mean-field approximation of synaptic dynamics for each neuronal population in the network (Wong and Wang, 2006; Deco et al., 2013; 2014b). Populations receive synaptic input from multiple sources, with contributions from the fluctuating background, local recurrent connections, and long-range connections from other areas, which induces structured correlated fluctuations across the network. Each local node is characterized by two synaptic parameters which set the strengths of local excitatory-to-excitatory (*w^EE^*) and excitatory-to-inhibitory (*w^EI^*) connections. The inhibitory-to-excitatory strength (*w^IE^*) was set to maintain a uniform baseline firing rate, dependent on the other parameters (Deco et al., 2014b). Global coupling parameters *g*_{*L*,*R*}_ scale the strengths of long-range interactions within the left and right hemispheres. Synaptic activity is used to simulate the BOLD signal using the mechanistic Balloon-Windkessel model for the hemodynamic response (Friston et al., 2003; Deco et al., 2013). We can thereby calculate a simulated BOLD FC matrix, which can be compared to empirical BOLD rs-FC data. The neurophysiological model parameters can then be optimized to provide the best fit to empirical rs-FC.

A key extension to the model framework introduced here is a hypothesis-driven approach to incorporate areal heterogeneity of local circuit properties (**Figure 1B**). We compared the performance of the circuit model with homogeneous and heterogeneous local circuit parameters. In the ‘homogeneous’ model, synaptic parameters were uniform across cortical areas, and its four parameters were optimized globally (*w^EE^*, *w^EI^*, *g_L_*, *g_R_*). In contrast, in the ‘heterogeneous’ model, fit parameter values — here, *w^EE^* and *w^EI^* — can vary across cortical areas, parametrized according to a pre-defined heterogeneity map. The heterogeneity map thereby constrains the topography of local circuit specialization in the heterogeneous model.

### T1w/T2w as a Hierarchical Heterogeneity Map

We hypothesized that cortical hierarchy provides a principle describing specialization of microcircuit properties across cortical areas which shapes large-scale functional dynamics. We therefore sought to implement the model with a heterogeneity map which reflects a hierarchical ordering of cortical areas. Because anatomical hierarchy is derived through invasive tract-tracing which has precluded direct investigation in human cortex, we sought a noninvasive proxy measure. Burt et al. (In press) found that the MRI-derived T1w/T2w map is negatively correlated with anatomical hierarchy in macaque cortex, and that specialization in multiple aspects of cortical microcircuitry were found to correlate with this T1w/T2w map. In particular, they found a negative correlation between T1w/T2w values and the number of spines on pyramidal cell dendrites. As the spine count can be interpreted as a microanatomical correlate for the strength of recurrent synaptic excitation in local cortical microcircuits (Elston, 2003; Chaudhuri et al., 2015), this finding suggests the model should incorporate stronger values of *w^EE^* in association areas with low T1w/T2w values (**Figure 1C**).

In human cortex, areas can be contextualized in terms of coherent resting-state networks (RSNs) associated with different sensory and higher-order association functions. We assigned all areas to 8 canonical RSNs comprising three sensory networks (visual, somatosensory, auditory) and five association networks (fronto-parietal, cingulo-opercular, default mode, dorsal attention, ventral attention) (Ito et al., 2017) (**Figure 1D**). We observed that T1w/T2w map values were significantly higher in sensory networks than in association networks (*p* < 0.003, Wilcoxon signed-rank test) (Burt et al., In press) (**Figure 1E**). In further support of the T1w/T2w map as a proxy measure of hierarchical microcircuit specialization across human cortex, Burt et al. (In press) analyzed the topography of cortical gene expression, and found that the T1w/T2w map captures the dominant spatial pattern of gene expression variation in human cortex. Of note, the T1w/T2w map outperformed the map of cortical thickness in capturing anatomical hierarchy in macaque and gene expression variation in human.

These findings suggest the T1w/T2w map may capture a key axis of areal heterogeneity across cortex, which we quantitatively instantiated in the model. We derived a hierarchical heterogeneity map by rescaling and inverting the raw T1w/T2w map, such that its values are relatively uniformly distributed between 0 and 1, with high-T1w/T2w sensory areas at low map values and low-T1w/T2w association areas at high map values. Local synaptic strengths (*w^EE^* and *w^EI^*) were then parametrized for each area *i* as an affine function of the heterogeneity map values {*h_i_*}, characterized by an intercept *w_min_* and scale factor *w_scale_*: *w_i_* = *w_min_* + *w_scale_h_i_* (**Figure 1B**). Use of a heterogeneity map in model fitting thereby enables hypothesis-driven investigation of areal differences in local circuit properties, while increasing model complexity by only a single additional parameter for each heterogeneous property.

### Model Fitting

We quantitatively fit the models described above to rs-FC data. To estimate the optimal model parameter values, we used hierarchical population Monte Carlo (hPMC), which is a Bayesian optimization technique (for details, see **Experimental Procedures**) (**Figure S3)**. HPMC approximates the posterior distribution in parameter space by iteratively drawing a set of model parameters (i.e., ‘particles’) from the proposed distribution to minimize a distance measure between model and empirical data. We fit the model parameters to maximize the average Pearson correlation between model and empirical FC across subjects (N=334). To calculate the model FC, we used an analytical approximation of linearized system dynamics, which enables computationally efficient calculation of dynamical features of the system, including the FC matrix (Deco et al., 2013; 2014b) (**Figure S7)**. Here we extended this approach to include linearization of the Balloon-Windkessel hemodynamic model for direct calculation of the BOLD FC matrix. The model fitting procedure produced to the approximated posterior distribution of the optimal model parameters for both homogeneous and heterogeneous models.

### Hierarchical Heterogeneity Improves Fit to FC

We tested whether introducing hierarchical heterogeneity improves the similarity between empirical rs-FC and the fit model FC patterns, compared to a homogeneous model with uniform local properties across cortical areas. **Figure 2A–C** shows the empirical group-averaged SC and FC matrices, and the particle-averaged FC matrices for the homogeneous and heterogeneous models. We quantified model performance using the fraction of explained variance (i.e., squared Pearson correlation coefficient) of the empirical FC, averaged across subjects, captured by the the model FC, averaged across samples from the approximate posterior distribution. We found that the similarity between empirical and model FC was significantly higher in the heterogeneous model (*r* = 0.563, *r*^2^ = 0.316) than in the homogeneous model (*r* = 0.407, *r*^2^ = 0.166) (*p* < 10^−4^, dependent correlation test). Compared to the SC-FC fit (*r* = 0.284, *r*^2^ = 0.081) as baseline, the model fits for both homogeneous and heterogeneous models were significantly larger (**Figure 2G**). The fit between SC and model FC was significantly lower for the heterogenous model (*r* = 0.413, *r*^2^ = 0.171) than the homogeneous model (*r* = 0.595, *r*^2^ = 0.354) (*p* < 10^−4^, dependent correlation test), which suggests that areal heterogeneity can help to explain FC patterns not accounted for by SC (Chaudhuri et al., 2015).

**Figure 2:**
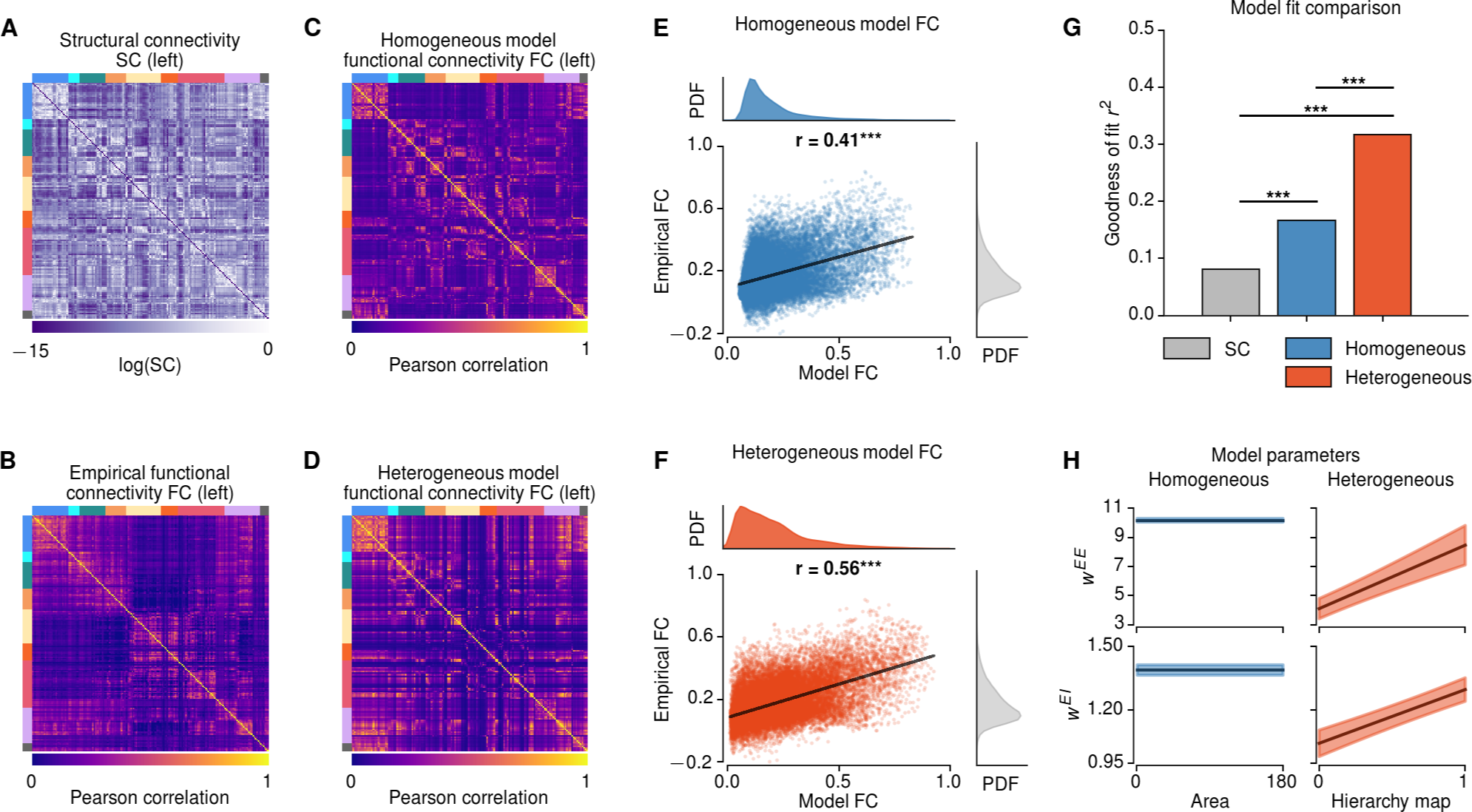
Hierarchical Heterogeneity Improves the Model Fit to Resting-State Functional Connectivity (rs-FC). **(A,B)** Structural connectivity (SC) and empirical functional connectivity (FC) matrices (left hemisphere only), averaged across subjects. Colored bars (top and left of matrices) denote resting-state network assignments (colored as in Fig. 1). **(C,D)** Model functional connectivity of the homogeneous and heterogeneous models (left hemisphere only), averaged across particles. **(E,F)** Correlation between average empirical FC and average model FC for the homogeneous and heterogeneous models. **(G)** Goodness of fit (i.e., fraction of explained variance *r*^2^) between the average empirical FC and the structural connectivity (gray), homogeneous model FC (blue), heterogeneous model FC (red). The fit for the heterogenous model is greater than that of the homogeneous model, which is greater than that of the structural connectivity (*p* < 10^−5^ for each, dependent correlation test). **(H)** The best-fit values for recurrent excitatory parameters for the homogeneous and heterogeneous models, with regions ordered by increasing values of the T1w/T2w-derived hierarchical heterogeneity map. Shaded regions show standard deviation across particles.

The optimal fit parameters for the heterogenous model exhibited a large scaling of local recurrent excitatory-to-excitatory synaptic strengths (*w^EE^*) across the hierarchical heterogeneity map 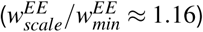 (**Figure 2H**). Interestingly, incorporation of heterogeneity expanded the dimensionality of fitting to the empirical population (**Figure S4)**. We applied principal component analysis (PCA) to the distribution of model particles from the hPMC fitting in the fit parameter space. The homogeneous model exhibited only a single axis of particle variation in its four-dimensional parameter space, indicating one-dimensional expressiveness of the synaptic parameters. In contrast, the heterogeneous model exhibited four dimensions of variation in its six-dimensional parameter space. The local recurrent excitatory-to-excitatory synaptic strength *w^EE^* played a prominent role in expanding this dimensionality (**Figure S4B–H**). These findings show that the heterogeneous model can improve the fit to empirical FC through increasing local recurrent strengths along the cortical hierarchy, with stronger connections in association regions relative to sensory regions, and that *w^EE^* has substantial influence on the model expressiveness (**Figure 2H**, **Figure S4B,G**).

Because the heterogeneous model has more parameters than the homogeneous model (6 vs. 4), we tested that the improved fitting was not due to over-fitting with the more expressive model. Implementing leave-p-out cross-validation, we repeated the fitting procedure for a randomly selected subset of 80% of the subjects (267), and measured the model fit with the remaining 20% of subjects (67). Across 100 cross-validation samples, the predictive power of the heterogeneous model (*r* = 0.55±0.0002) always outperformed the homogeneous model (*r* = 0.40±0.0002).

In addition, we tested whether the improved model fit in heterogeneous model can be explained by known non-neural confounds in rs-fMRI, such as head motion, variations in heart rate, and respiration. If the improved fit of the heterogeneous model were due to its capture of non-neural FC contributions, then individual differences in non-neural measures should explain individual differences in the improvement in empirical-model FC similarity for heterogeneous vs. homogenous models. Across the 334 subjects, we found that the heterogeneous model improved the FC fit for 98.5% of subjects. We performed a regression analysis across subjects on the difference in model-empirical fit between heterogeneous and homogeneous models. A constant term (i.e., without non-neural measures) explained 80% of the total sum of squares, and inclusion of individual non-neural measures improved the explained variance by only 1.5%. This indicates that the substantial improvement in fitting by the heterogeneous model is not attributable to non-neural confounds.

### T1w/T2w Map as Preferential Axis of Areal Specialization

Does the T1w/T2w-derived hierarchical heterogeneity map provide a preferential axis of cortical specialization, in comparison to other possible heterogeneity maps? To address this question, we repeated the model fitting procedure for a single hemisphere using the T1w/T2w-derived heterogeneity map and 500 randomized surrogate heterogeneity maps. Because T1w/T2w map values are spatially autocorrelated, we developed a procedure to generate surrogate maps which randomly vary in their particular topographies but preserve the general spatial autocorrelation structure of the T1w/T2w-derived map (see **Experimental Procedures**) (**Figure 3A–C**).

**Figure 3:**
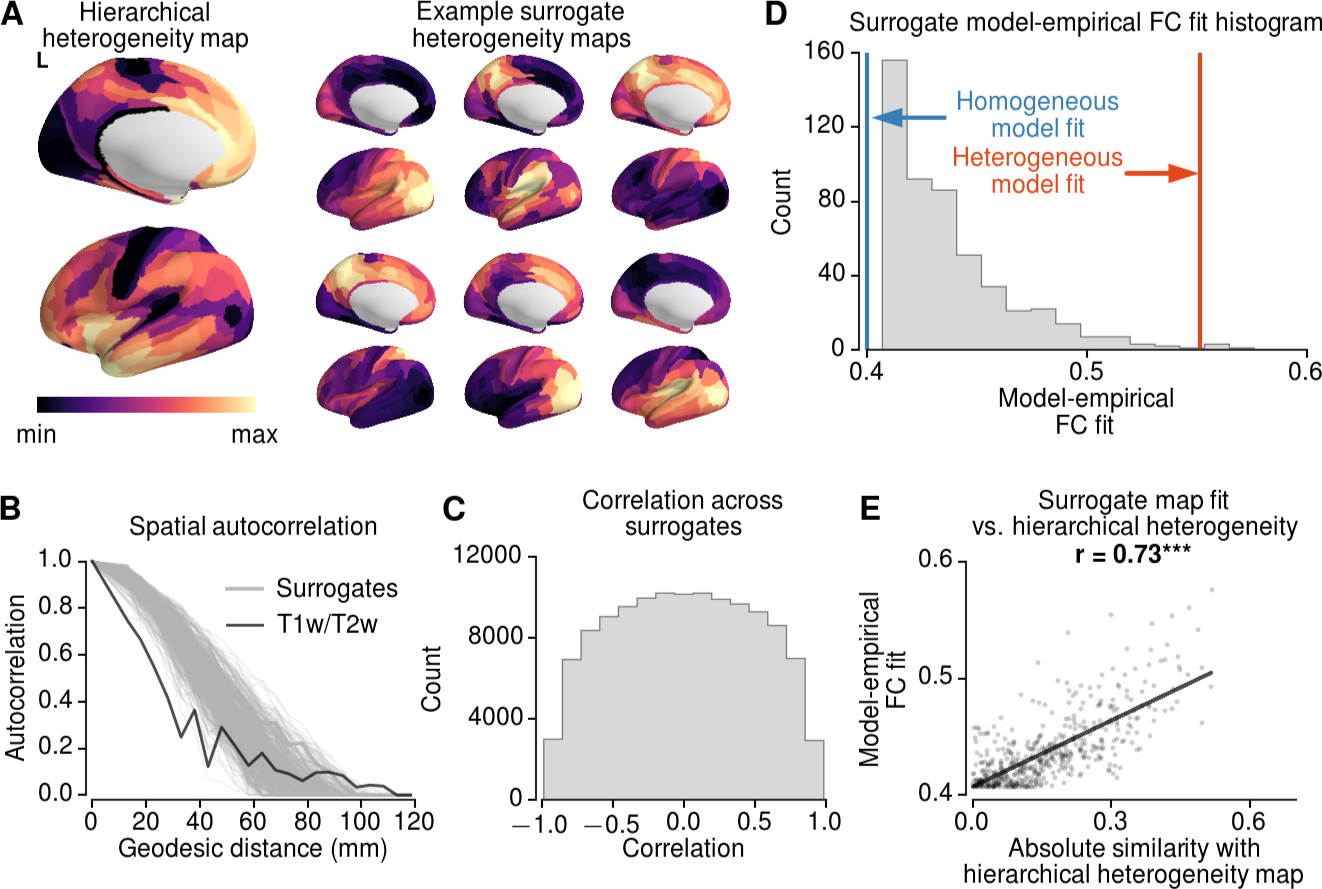
Surrogate Heterogeneity Maps Show the T1w/T2w Map Provides a Preferential Axis of Specialization. **(A)** The T1w/T2w-derived hierarchical heterogeneity map, for the left hemisphere, and example surrogate heterogeneity maps with matched spatial autocorrelations. **(B)** Spatial autocorrelations of Box-Cox-transformed T1w/T2w map (black) and surrogate heterogeneity maps (gray) as a function of geodesic distance. **(C)** Histogram of spatial correlations (Spearman rank) between all pairs of random surrogate maps. **(D)** Histogram of the best fit (correlation between empirical and model FC) of random surrogates. The T1w/T2w map gradient fit is significantly higher than random surrogates (*p* = 0.008). **(E)** The correlation between hierarchical heterogeneity-surrogate map similarity (i.e., absolute values of correlation) and model performance (i.e., model-empirical FC similarity). The model-empirical FC similarities for surrogate maps increase with the absolute value of the correlation with hierarchical heterogeneity map.

We found that the model fit of the surrogate maps were slightly higher than that of the homogeneous model, with an average correlation *r* = 0.43 for surrogates vs. *r* = 0.40 for the homogeneous model. Nonetheless, among all surrogate maps, the T1w/T2w-derive map exhibited significantly higher model fit than surrogates (*p* = 0.008, *r* = 0.55 for the heterogenous model) (**Figure 3D**). Furthermore, the model fit using surrogate heterogeneity maps was were significantly correlated with their similarity to the T1w/T2w-derived heterogeneity map (**Figure 3E**). Therefore, the substantial improvement in the model performance was strongly preferential to the T1w/T2w-derived hierarchical heterogeneity map. These findings suggest that the T1w/T2w map provides a preferential neural axis for cortical microcircuit specialization, in line with prior empirical characterization of microanatomical and transcriptional specialization along the T1w/T2w map (Burt et al., In press).

### Model Fit in Resting-State Networks

We examined how the improved performance of the hierarchical heterogeneous model was distributed across different functionally relevant cortical networks, and whether this was due to capturing rs-FC structure within networks or across networks. We calculated the model-empirical FC similarity for each of eight RSNs (three sensory, five association), decomposing its FC pattern into within-network (**Figure 4A**) and across-network (**Figure 4D**) components. We found that both within- and across-network fits to empirical FC were higher in the heterogeneous model than in the homogeneous model for all RSNs (**Figure 4B–F**), with association RSNs showing a larger improvement than sensory RSNs in within-network similarity. Among the association RSNs, the frontoparietal (FPN) and cingulo-opercular (CON) networks exhibited large increases in both within- and across-network fits. These results show that incorporating hierarchical heterogeneity not only improves the whole-cortex model FC fit, it also preferentially improves the within-network model FC fit in association networks.

**Figure 4:**
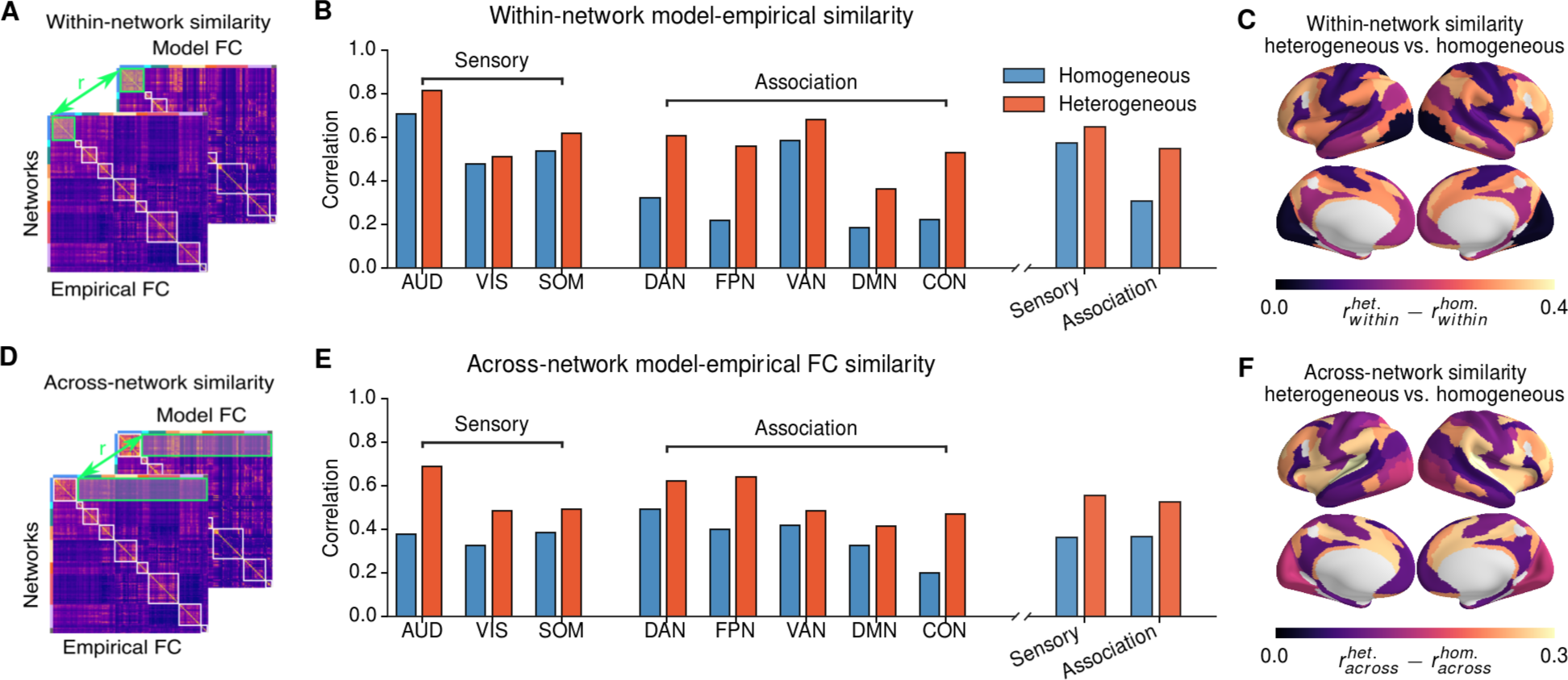
Model Fits Across Resting-State Networks (RSNs) are Network-Specific. **(A,D)** Schematic of within- and across-network fits. Correlations between empirical and model FC within or between RSNs were calculated for homogeneous and heterogeneous models. **(B,E)** Within- and across-network fits of the models. Heterogeneous model showed substantial improvements compared to homogeneous model for within- and across-networks fit in all networks. Across-network fit improvements were distributed across sensory and association networks. Within-network fit improvements were preferentially in association networks. **(C,F)** Topography of the improvement in fit for heterogeneous vs. homogeneous models. Values are shown for each RSN.

### Global Brain Connectivity

The RSN analyses presented above suggest that hierarchical heterogeneity may allow the model to capture important rs-FC differences between sensory and association networks. To investigate sensory–association differences more directly, we studied the topography of global brain connectivity (GBC), a measure of global FC strength for each area (**Figure 5A**). Multiple studies have found GBC to be an informative measure of rs-FC alterations in psychiatric disorders (Cole et al., 2011; Yang et al., 2016; Demirtaş et al., 2017) and individual differences in cognition (Cole et al., 2012). Furthermore, there is evidence that the cortical topography of pharmacologically-induced changes in GBC is aligned with the topography of gene expression for its targeted receptor (Preller et al., 2017). We therefore examined whether hierarchical heterogeneity of circuit properties in the model shapes the cortical GBC topography in healthy subjects.

**Figure 5:**
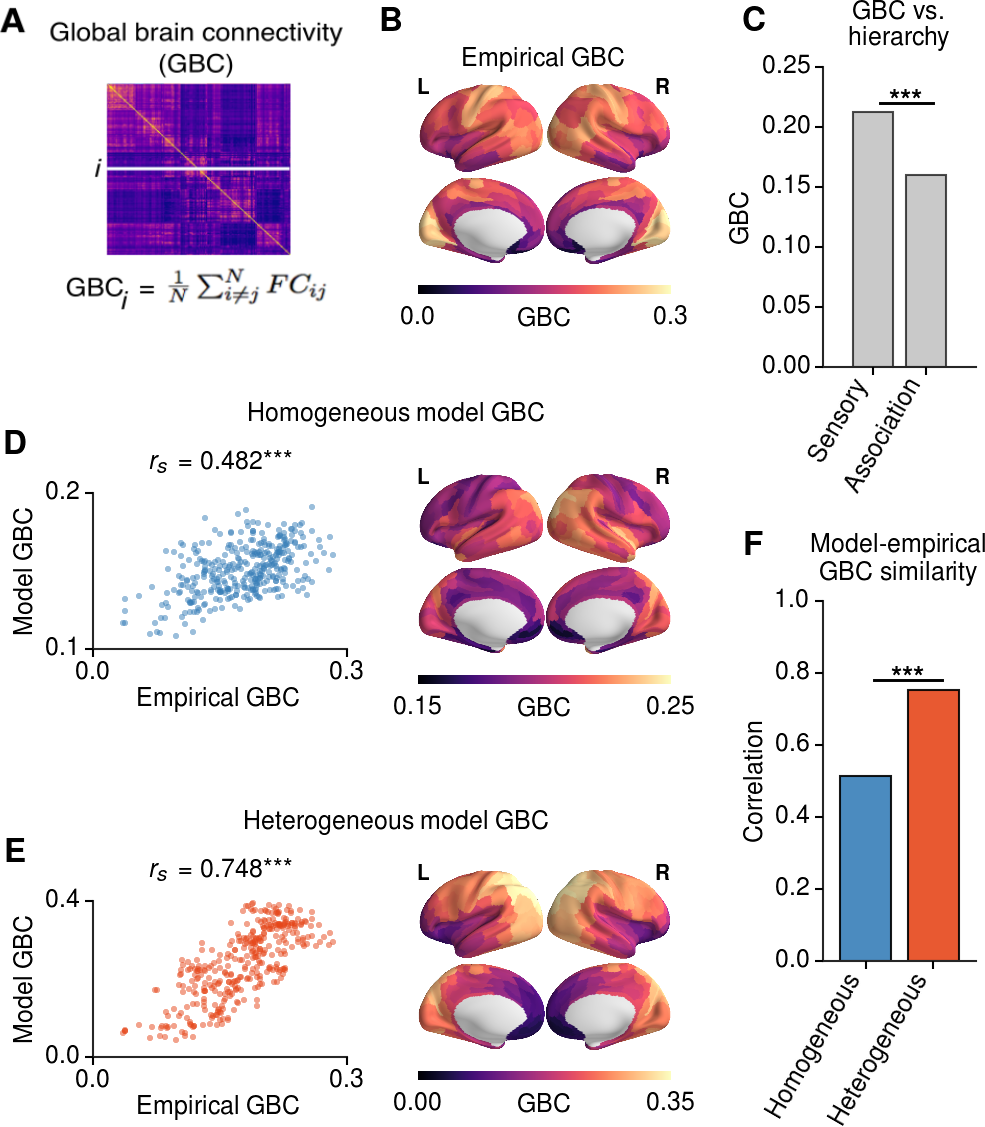
Hierarchical Topography of Cortical Global Brain Connectivity (GBC). **(A)** GBC of each region is calculated as the average FC of that region with all other cortical regions. **(B)** The areal topography of empirical GBC. **(C)** GBC of sensory areas is significantly larger than that of association areas (*p* < 0.001, Wilcoxon signed-rank test). **(D–F)** The correlation between empirical and model GBC is significantly larger in the heterogeneous model than in the homogeneous model (*p* < 10^−5^, dependent correlation test).

We found that GBC, calculated within cortex, was significantly different across sensory and association RSNs, with higher GBC in higher GBC in sensory areas than in association areas (*p* < 0.003, Wilcoxon signed-rank test) (**Figure 5C**). The heterogeneous model the correlation with empirical GBC (*r_s_* = 0.749) was significantly higher than that in the homogeneous model (*r_s_* = 0.482) (*p* < 10^−4^, dependent correlation test) (**Figure 5D,E**). These findings suggest that hierarchical heterogeneity of local circuit properties shapes sensory–association differences in the large-scale organization of rs-FC. Furthermore, these findings suggest that observations of GBC alterations, and their topographies, may be driven by differences in local circuit properties.

### Inter-Individual Variation

Cortical rs-FC patterns vary across individuals. Mueller et al. (2013) characterized the degree to which cortical areas vary in their FC profiles across subjects, and found a marked hierarchical difference across RSNs: sensorimotor regions exhibited low inter-individual variation in FC, whereas association regions exhibited higher variation. As noted above, our model fitting approach uses hPMC to fit a posterior distribution in parameter space to the full set of FC patterns across subjects in the HCP dataset. Of note, particles share the same structural connectivity matrix and heterogeneity map, and differ only in their synaptic parameter values. We can therefore study whether the best-fit model exhibits hierarchical differences in FC variation across particles drawn from the approximated posterior distribution, comparable to the FC variation across subjects in the empirical data.

We quantified the variability within the population as the dissimilarity of FC patterns across individuals, following the approach of Mueller et al. (2013). In the model, the dissimilarity of FC patterns was calculated across 1,000 particles that were sampled from the posterior distribution (**Figure 6A**). In this study, model particles all used the same SC matrix, averaged over subjects. This allowed our analyses to isolate potential contributions to individual variation in rs-FC arising from variation in physiological circuit properties. We note this is one potential source of variation, and that individual differences in SC likely also shape differences in FC (Zimmermann et al., 2018).

**Figure 6:**
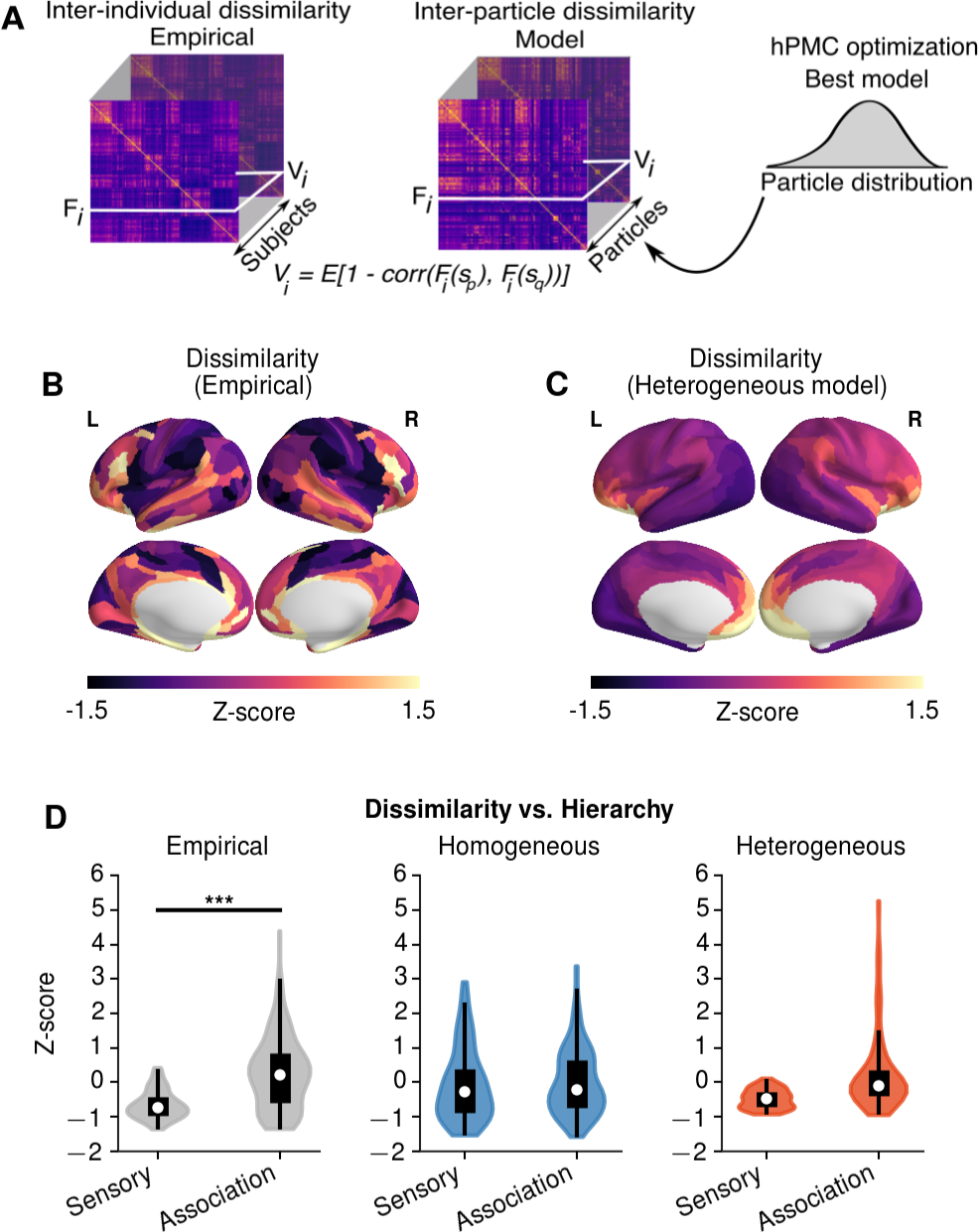
Hierarchical Topography of Inter-Individual Dissimilarity of FC. **(A)** Dissimilarity calculated for FC patterns across subjects (*N* = 334) in the empirical dataset, and across particles (*N* = 1000) in the model fitting framework. The dissimilarity for area *i*, *V_i_*, is given by *V_i_* = E(1 − Corr(*F_i_*(*s_p_*),*F_i_*(*s_q_*)), where *E*(…) is the mean across subject pairs, and *F_i_*(*s_p_*) is the FC of area *i* for subject *s_p_*. To compare the regional topography in empirical and model dissimilarity maps, we standardize the values through z-score. **(B-C)** Topography of empirical inter-individual dissimilarity and heterogeneous model inter-particle dissimilarity. The inter-particle dissimilarity for the homogeneous model was not depicted due to lack of spatial patterns. **(D)** Inter-individual dissimilarity is higher for association areas than than sensory areas (*p* < 0.003, Wilcoxon signed-rank test). The heterogeneous model exhibits a similar hierarchical differentiation in inter-particle dissimilarity. The association–sensory difference is larger in the heterogeneous model than homogeneous (*p* < 0.0001, 2 × 2 ANOVA, *z* = 4.4175).

Across 334 HCP subjects, we found that the inter-individual dissimilarity was higher in frontal and temporal brain regions(**Figure 6B**). The inter-individual dissimilarity in association RSNs (DAN, VAN, FPN, DMN, CON) was on average higher than in sensory RSNs (VIS, AUD, SOM) (*p* < 0.003, Wilcoxon signed-rank test; z-score 0.265±1.055 and –0.659±0.410 for association and sensory RSNs, respectively) (**Figure 6D**). Similar to the empirical data, the heterogeneous model showed higher inter-particle dissimilarity in frontal association areas (**Figure 6C**). The topography of empirical dissimilarity was positively correlated with that of the heterogeneous model (*r* = 0.51), unlike with the homogeneous model (*r* = –0.05). The heterogeneous model exhibits a similar hierarchical distinction in dissimilarity, with higher inter-particle variability in association regions than in sensory regions, whereas no such pattern is present in the homogeneous model (*p* < 0.0001, network × model type ANOVA, *z* = 4.4175) (**Figure 6D**). These findings suggest hierarchical heterogeneity may contribute to individual variation in the functional organization of cortex.

### Heterogeneity in Neural Dynamics Across Multiple Timescales

How does hierarchical heterogeneity in the model shape neural dynamics across a wide range of timescales? Results described above examined simulated BOLD signals from a hemodynamic model which are driven by synaptic activity in the neural circuits. Cortical areas differ in the spectral properties of their intrinsic dynamics at rest (Honey et al., 2012; Murray et al., 2014; Keitel and Gross, 2016; Mellem et al., 2017). In local circuit models, synaptic strengths can shape these spectral properties (Chaudhuri et al., 2015; Murray et al., 2017). We therefore sought to examine in the model how hierarchical heterogeneity produces specific areal topographies of neural dynamics across multiple timescales.

We first characterized the dynamical repertoire of the local microcircuit model at each node in the large-scale network, as a function of the recurrent excitatory strengths onto excitatory and inhibitory neurons: *w^EE^* and *w^EI^* (**Figure 7A**, **Figure S5)**. As *w^EE^* increases, there is a threshold value, i.e., a bifurcation point, beyond which the system’s baseline state destabilizes (Deco et al., 2013). When *w^EI^* is large, the system exhibits another dynamical regime in which the system’s dynamics undergo damped oscillations (**Figure 7A**). In this oscillatory regime, the intrinsic frequency of depends on *w^EE^* and *w^EI^* (**Figure 7A**). Therefore, in the higher-dimensional parameter space of microcircuits in the heterogeneous model, the model can enter more dynamical regimes of asynchronous and damped oscillatory dynamics depending on the local excitation-inhibition balance. We examined the optimal parameter ranges of homogeneous and heterogeneous models. We found that the optimal solutions of homogeneous and heterogeneous models were close to the critical points, in line with prior studies (Deco et al., 2013; 2014b). The optimal solutions of the homogeneous model were found where each node exhibited oscillatory dynamics (**Figure 7B**), whereas in the heterogeneous model the dynamical range of the nodes spanned asynchronous and oscillatory dynamics (**Figure 7C**).

**Figure 7:**
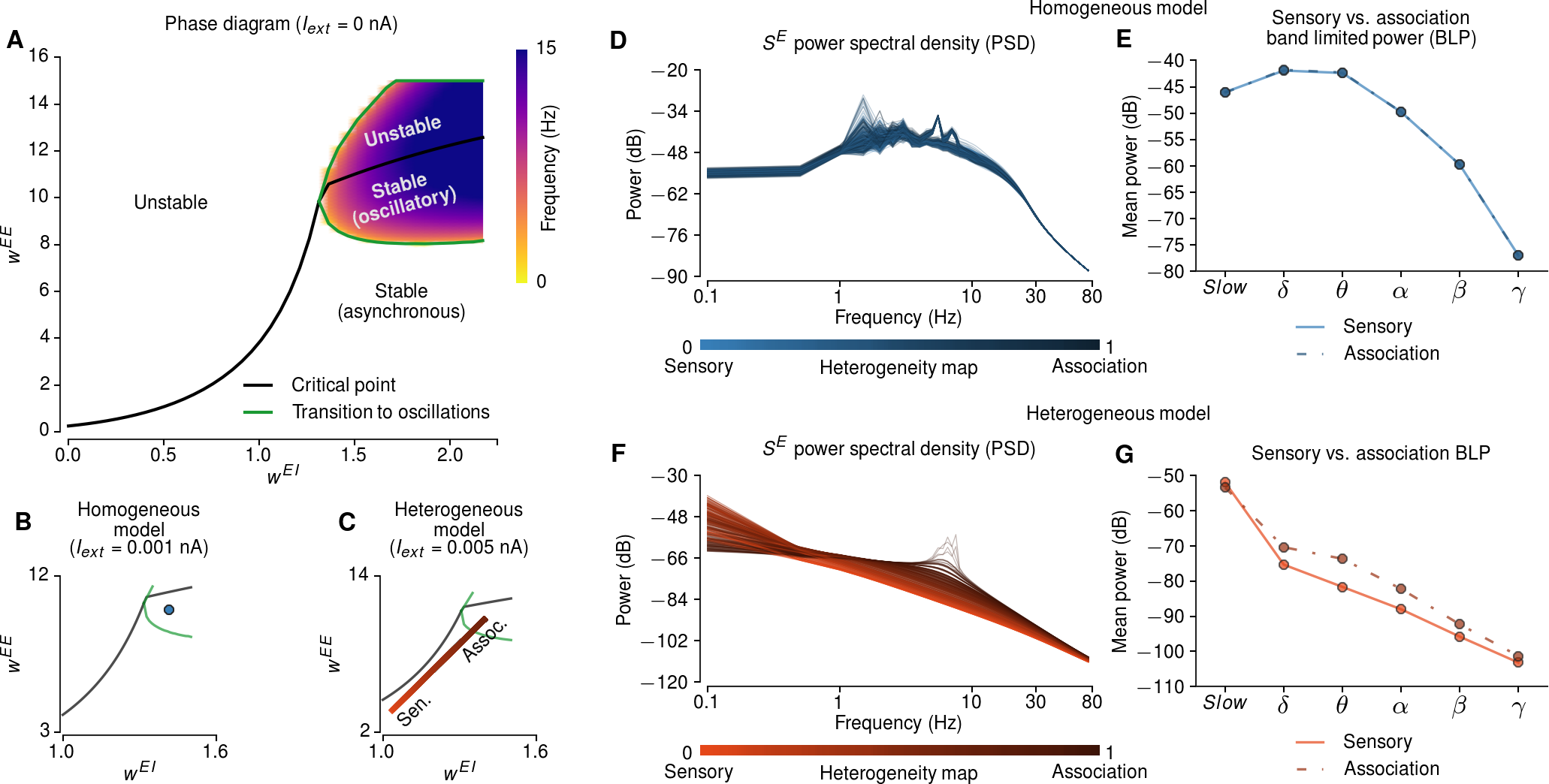
Intrinsic Dynamics of a Local Microcircuit Model Vary with Recurrent Strengths. **(A)** The phase diagram of a local microcircuit model (i.e., one node in the large-scale network) without external input. The black line indicates the critical points beyond which the baseline state is unstable, and the green lines indicate the boundaries at which the system exhibits a transition to oscillatory dynamics. For excitatory-to-inhibitory synaptic strengths (*w^EI^*) smaller than 1.35 the system exhibits a pitchfork bifurcation (in asynchronous dynamics), and for larger *w^EI^* values the system exhibits a transition to damped oscillatory dynamics. **(B,C)** A representative example of optimal homogeneous and heterogeneous model parameters projected onto the phase diagram. The external input is adjusted to provide the same mean long-range input as in the fit large-scale model from other areas. **(D,F)** The power spectral densities (PSDs) of the synaptic gating variables (*S^E^*) of the homogeneous (**D**) and heterogeneous (**F**) models. The colors indicate the hierarchical level of each area based on its T1w/T2w map value (light: sensory, dark: association). **(E,G)** The relative power of sensory and association regions of homogeneous (**E**) and heterogeneous (**G**) models in discrete frequency bands: slow (<2 Hz), delta (*δ*; 2–4 Hz), theta (*θ*; 4–8 Hz), alpha (*α*; 8–15 Hz), beta (*β*; 15–35 Hz) and gamma (*γ*; 35–50 Hz).

We characterized the power spectral properties of the underlying synaptic activity in the heterogeneous and homogeneous models. We calculated the power spectral density (PSD) across parcels in frequencies ranging between 0 and 80 Hz. To investigate the contrast between sensory and association regions, we computed the mean band limited power (BLP) of sensory and association regions in five commonly-used frequency bands: slow (< 2 Hz), delta (2–4 Hz), theta (4–8 Hz), alpha (8–15 Hz), beta (15–35 Hz), and gamma (35–50 Hz) bands. In the homogeneous model, the contribution of sensory and association regions to BLPs did not differ substantially (**Figure 7D–E**). In contrast, the heterogeneous model exhibited PSDs broadly distributed across frequencies. At slow frequencies (<2 Hz) the BLP was higher in sensory areas, whereas in faster frequencies the BLP was higher in association areas peaking in the theta range (2–4 Hz) (**Figure 7F–G**). To examine the contributions of inter-areal connectivity to these dynamics, we simulated the effect of “disconnection” in the network (**Figure S6)**. We found that long-range connections primarily shape low-frequency power (< 1 Hz), suggesting differential contribution of local and long-range inputs in shaping spectral properties. Furthermore, we found that introducing distance-dependent synaptic delays in long-range projections did not substantially affect simulated BLPs or BOLD FC patterns (**Figure S6)**. These results suggest that hierarchical heterogeneity predicts specific spatial distribution of spectral power across multiple timescales.

### MEG Power Spectral Density

We tested the prediction that spectral patterns are hierarchically organized using the PSD from resting-state MEG averaged across subjects (**Figure 8A**). The empirical MEG-derived PSD of each parcel was normalized across frequencies and relative band limited power (BLP) maps were calculated for 5 frequency bands. We found that the raw T1w/T2w-derived hierarchical heterogeneity map is substantially correlated with BLP spatial patterns in all frequency bands (**Figure 8B**). For each frequency band, we calculated the significance values based on a spatial lag model which accounts for spatial autocorrelation in the maps.

**Figure 8:**
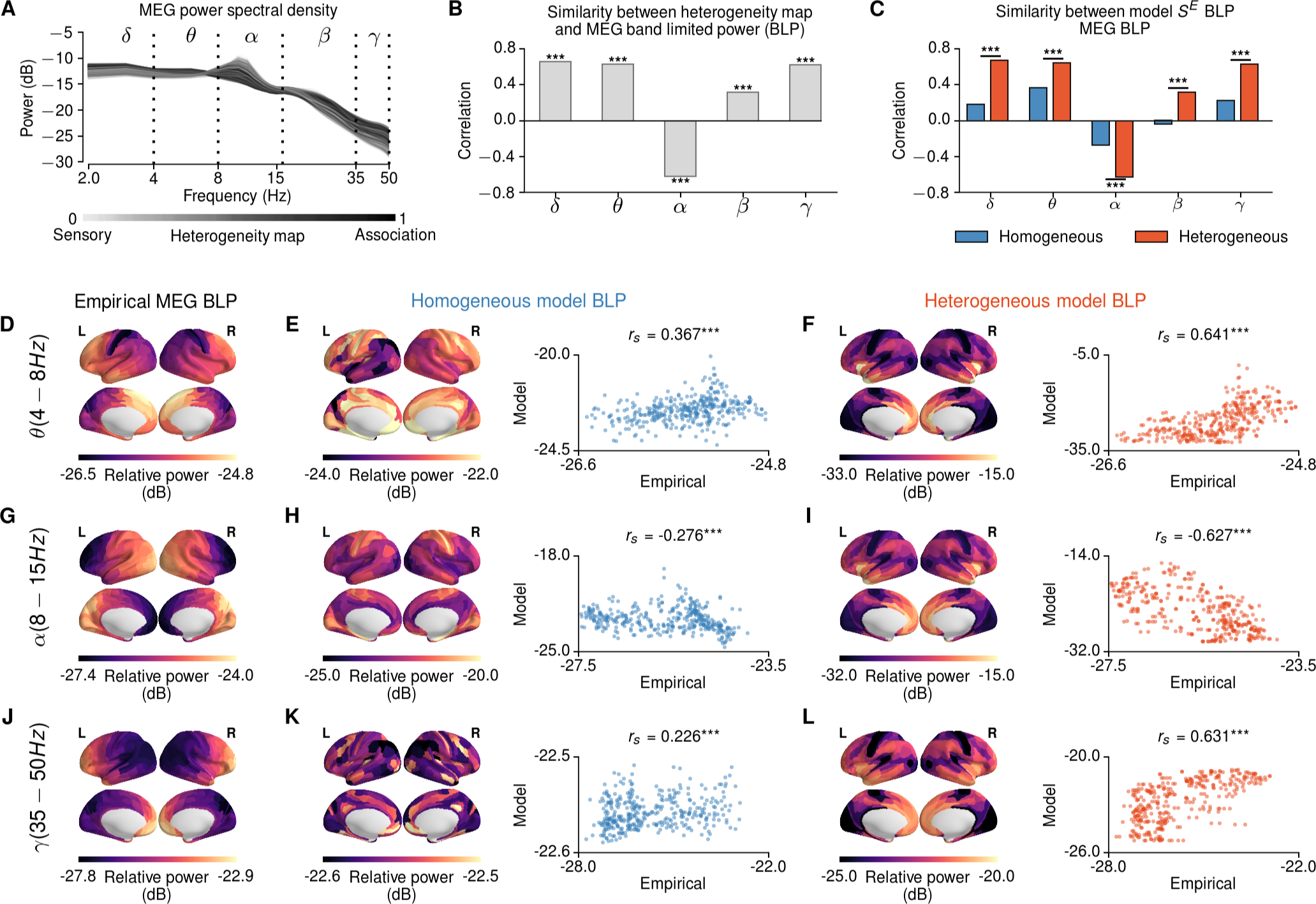
Hierarchical Topography of Spectral Power in Magnetoencephalograpy (MEG). **(A)** The empirical PSD derived from MEG. The shading of lines indicate the hierarchical level of each region based on its T1w/T2w map value (light: high (sensory), dark: low (association)). Dashed lines mark commonly defined frequency bands: delta (*δ*; 2–4 Hz), theta (*θ*; 4–8 Hz), alpha (α; 8–15 Hz), beta (*β*; 15–35 Hz) and gamma (*γ*; 35–50 Hz). **(B)** Correlations between the maps of MEG band-limited power (BLP) and the T1w/T2w-derived hierarchical heterogeneity map. The significance of the correlations were calculated using a spatial lag model, FDR-corrected for multiple comparisons. *** denotes *p* < 0.001, * denotes *p* < 0.05, n.s. denotes *p* > 0.05. **(C)** Correlations between MEG and model BLP maps. The magnitudes of correlation with empirical MEG maps are higher for the heterogeneous model (*p* < 10^−5^, dependent correlation test). **(D–L)** shows the relationship between empirical and heterogeneous model BLPs in each frequency band.

The correlations between empirical and model BLPs were significantly higher for the heterogeneous model than the homogeneous model (*p* < 10^−5^, dependent correlation test) (**Figure 8C**). The empirical-model BLP correlations were highest in the delta (*r_s_* = 0.67), theta (*r_s_* = 0.64) and gamma (*r_s_* = 0.63) bands (**Figure 8E**, **Figure S8)**. The BLP correlation was significantly negative in the alpha band (*r_s_* = –0.627), which has been shown to involve subcortical sources such as coupling between thalamus and occipital cortex (Hughes and Crunelli, 2005). The correlations between MEG BLPs and homogeneous model BLPs were weak and significantly lower than those in the heterogeneous model. These results show that the BLPs in MEG also show topographic signatures of hierarchical heterogeneity across cortical areas.

## Discussion

In this study, we proposed a biophysically-based large-scale dynamical model of human cortical activity that incorporates hierarchical heterogeneity in local circuit properties. Specifically, we parametrized local excitatory synaptic strengths following the topography of the T1w/T2w map. Incorporating hierarchical heterogeneity substantially improved the model fit to BOLD rs-FC and captured thesensory-association organization of multiple fMRI features. The heterogeneous model predicted spatially patterned spectral properties across a wide range of timescales, which were consistent with MEG. Collectively, our findings suggest that heterogeneity in local circuit properties of shapes the large-scale organization of neural dynamics in human cortex.

We hypothesized that the T1w/T2w map would provide a key neural axis along which microcircuit properties and spatiotemporal dynamics would vary. This hypothesis was informed by a convergence of findings across multiple modalities linking the T1w/T2w map, microcircuit specialization, functional organization, and the organizing principle of cortical hierarchy.

The T1w/T2w map captures a number of aspects of microcircuit specialization across cortex. This structural MRI-derived contrast measure is sensitive to regional variation in multiple microstructural properties including intracortical myelin content (Glasser et al., 2014; 2016). Burt et al. (In press) found that the T1w/T2w map captures areal variation in cytoarchitecture (Hilgetag et al., 2016), cell type distributions, and synaptic properties. The number and density of dendritic spines on pyramidal neurons, a microanatomical correlate of local synaptic excitation, increase with cortical hierarchy as captured by the T1w/T2w map (Elston, 2003; Glasser et al., 2014; Chaudhuri et al., 2015; Burt et al., In press). This observation provides microanatomical support for our heterogeneous model’s local recurrent excitatory strengths following an increasing hierarchical gradient (Chaudhuri et al., 2015). Furthermore, large-scale gene expression mapping of human cortex reveals that the T1w/T2w map captures the dominant neural axis of transcriptional variation, suggesting a common hierarchical organization for functional specialization (Burt et al., In press). These findings support our use of the T1w/T2w map as a key neural axis for hierarchical microcircuit specialization.

Electrophysiological studies have established that functional dynamics of cortical areas also exhibit an organization along a sensory-association hierarchy. The timescales of intrinsic fluctuations vary across hierarchical levels both in humans (Honey et al., 2012) and in monkeys at the single-neuron level (Murray et al., 2014). Computational modeling studies have demonstrated that these differences can arise from hierarchical differences in synaptic properties, including excitatory strengths. In turn, models have shown how hierarchical differences in synaptic properties can contribute to functional specialization of areas, such as their capacity to generate the robust persistent activity thought to underlie working memory computations (Wang, 2001; Chaudhuri et al., 2015; Murray et al., 2017).

FMRI has also revealed functional organization of human cortex along sensory-association hierarchical gradients. For instance, Margulies et al. (2016) found that the principal gradient of rs-FC variation across cortex separated primary sensory areas from higher-order association areas. Furthermore, this gradient aligns with the topography of intracortical myelin content, as measured by T1-mapping (Huntenburg et al., 2017). Compared to sensory cortex, association areas exhibit greater variation in rs-FC profiles across subjects, suggesting that cortical hierarchy plays a role in shaping single-subject specificity of FC patterns (Mueller et al., 2013; Finn et al., 2015). Our model provides a circuit mechanism linking hierarchical variation in microcircuit properties to functional specialization.

The association between dMRI-derived SC and rs-FC in human cortex has been consistently shown in prior studies (Greicius et al., 2009; Hagmann et al., 2008; Honey et al., 2009; Hermundstad et al., 2013; Baria et al., 2013), and provides the foundation for large-scale structure-function relationships. Computational studies of large-scale cortical circuits have shown that the structure-function relationship improves when neural dynamics are near the edge of instability (Deco et al., 2013; 2014b; 2017), which was also confirmed in our study. Despite the substantial performance of computational models in predicting rs-FC, the contribution of local neural circuit specialization has not characterized. Here we showed that the heterogeneity in local microcircuit properties has a substantial effect on model FC predictions, beyond prior modeling studies. In addition to the role of SC in shaping FC, this study provides a different, complementary perspective that emphasizes the importance of the intrinsic properties within local circuits in shaping the large-scale functional organization of human cortex.

Beyond the T1w/T2w-derived hierarchical heterogeneity map examined in this study, our model framework can be flexibly extended to include other axes of variation. For instance, heterogeneity maps could reflect varying distributions of receptor subtypes, neuronal cell types, and neuromodulators. These maps could be derived from gene expression (Burt et al., In press), positron emission tomography (Beliveau et al., 2017), or autoradiography (Palomero-Gallagher and Zilles, 2017). This direction has the potential to allow large-scale models to be fitted to pharmacological neuroimaging, through simulation of hypothesized parameter perturbations which follow the expression topography of the affected receptors (Preller et al., 2017). This can further be applied to clinical neuroimaging data, to fit the heterogeneous effects of disorder-related alterations (Yang et al., 2016). Further computational studies of areal heterogeneity can further inform the relationships between structure, function, and physiology in the human brain.

An important limitation in modeling neural mechanisms underlying rs-FC is the existence of multiple, potentially confounding, non-neural contributions to noninvasive neuroimaging measures, such as head motion, respiration and heart-rate variability (Power et al., 2017; Siegel et al., 2017). We addressed these issues by demonstrating that individual differences in non-neural measures did not explain the improvement in model-empirical FC fit by the heterogeneous model. Furthermore, the model predictions at faster timescales are consistent with spectral patterns that are empirically observed in MEG, which relies on electrophysiology rather than BOLD. Nevertheless, complete removal of, or control for, non-neural confounds in a given neuroimaging modality is difficult. Characterization of neural sources of rs-FC components can be provided through convergence of multiple modalities (Kucyi et al., 2018), including invasive recordings in animal models (Schölvinck et al., 2010; Mateo et al., 2017).

Our heterogeneous circuit model is parsimonious, as it simulates simplified local node dynamics, comprises only cortex, and permits a single neural axis of microcircuit specialization. The model can be extended in multiple important directions, which will expand the range of questions it can address in future studies. The local node model used here consists of a single excitatory and inhibitory population per area, which is an appropriate scale for the limited resolution of fMRI. Beyond synaptic properties considered in our study, one aspect of microcircuitry which exhibits hierarchical gradients is the distributions of multiple classes of inhibitory interneurons (Burt et al., In press), which differentially shape circuit dynamics and function (Womelsdorf et al., 2014).

Another important future direction will be to study an extended cortical microcircuit model with multiple layers. Superficial and deep layers are differentially involved in the oscillatory dynamics of spontaneous cortical activity (Mejias et al., 2016). This extension may therefore enable better fitting to the PSD across a wide range of timescales. Furthermore, a laminar model will allow for layer-specific long-range projections which exhibit a hierarchical organization in feedforward and feedback signaling (Michalareas et al., 2016). The richer temporal dynamics likely cannot be resolved by fMRI but can potentially be constrained by MEG, EEG, or ECoG. Multimodal functional neuroimaging may play a key role in development of biophysically based circuit models that operate across a wide range of spatiotemporal scales.

Another promising direction for model extension is to incorporate subcortical structures. For instance, our cortical model does not capture the strong alpha-band power in occipital cortex, which is thought to originate from interactions between cortex and thalamus (Hughes and Crunelli, 2005). Dynamical neural models of a thalamo-cortical loop can capture key aspects of resting-state dynamics, such as the temporal statistics of alpha-band activity in occipital cortex measured with EEG (Freyer et al., 2009; 2011). Future large-scale modeling can be extended to include multiple thalamic nuclei, with distinct bi-directional interactions with cortex. Our circuit modeling framework is well suited to study how large-scale recurrent cortico-subcortical interactions shape the spatiotemporal dynamics of cortex.

In conclusion, we report a large-scale neural circuit model of human cortex with hierarchical heterogeneity of local microcircuit properties across cortical areas. The model proposes a specific circuit mechanism for how microcircuit specialization shapes the large-scale functional organization of human cortical dynamics. Our findings highlight the importance of regional heterogeneity in local circuit properties, and provide support for a hierarchical neural axis reflecting important structure-function relationships in the human brain.

## Experimental Procedures

### Multi-modal Neuroimaging Dataset

We used resting-state fMRI (rs-fMRI) and diffusion MRI (dMRI) of 334 unrelated subjects from the Human Connectome Project (HCP) 900-subject data release (12/08/2015) (Van Essen et al., 2013). The magnetic resonance (MR) preprocessing relied on the surface-based multimodal intersubject registration (MSMAll) (Robinson et al., 2014). The analyses also involved resting-state magnetoencephalography (MEG) data of 89 subjects from the same HCP dataset. The analyses were focused on the cerebral cortex, and areas were defined according to a multimodal parcellation (MMP1.0) comprising 360 areas, 180 per hemsiphere, using the 210P boundaries from Glasser et al. (2016).

### Resting-State Functional Connectivity

The preprocessing of rs-fMRI time series were done according to the HCP minimal preprocessing pipeline (Glasser et al., 2013). BOLD time series were further denoised using FIX-ICA which yielded the signal that drove the cortical parcellation used in this study (Glasser et al., 2016). The rs-fMRI time series of each subject comprised 4 sessions each spanning 15 minutes recorded with repetition time (TR) 0.72 s. The rs-fMRI time series were parcellated into 360 areas (180 areas per hemisphere) using the MMP1.0 parcellation (Glasser et al., 2016). No further preprocessing step was performed to ensure that the time series are consistent with those produced the parcellations as referenced in Glasser et al. (2016).

We removed the first 100 time points from each of the BOLD scans to mitigate for any baseline offsets or signal intensity variation. In turned, we z-scored the time series of each area, the parcellated time series of each subject were concatenated to a single time series comprising 4400 time points (52.8 minutes). The time series of each area was z-scored again after concatenation. The functional connectivity (FC) matrix of each subject was computed using Pearson’s correlation coefficient between the time series of all pairs of areas.

### Structural Connectivity

Structural connectivity (SC) matrices were constructed using probabilistic tractography, for each of the 334 subjects, from the HCP diffusion MRI (dMRI) minimally preprocessed data (Glasser et al., 2013; Sotiropoulos et al., 2013). Following the HCP dMRI step, the diffusion images were underwent FSL’s bedpostx and probtrackx2 analysis workflows for probabilistic tractography. The SC matrices were derived by seeding at the white matter-gray matter boundary interface and counting the number of streamlines that intersected 60,000 white matter-gray matter boundary locations. Fiber orientations used in tractography were derived using a parametric deconvolution approach available in FSL (Jbabdi et al., 2012; Hernández et al., 2013; Sotiropoulos et al., 2016) (up to 3 orientations per voxel).

The dense connectome was then parcellated by considering the average between pairs of areas, and the resulting SC matrices were averaged across subjects. The diagonal elements of the group-averaged SC matrix were removed and then values of the SC was normalized between 0 and 1 (i.e., the SC matrix was divided by the maximum SC weight). For use in the computational model, the SC matrix was row-normalized, so that the summed long-range input strengths are equalized for each node in the network, which is simulated as a cortical microcircuit.

### T1w/T2w maps

The MSMAll registered and bias field corrected maps of the ratio between T1- to T2-weighted images (T1w/T2w) were provided with HCP dataset. In the MMP parcellation, each of the 180 areas in each hemisphere is assigned a paired homologues in the other hemisphere (Glasser et al., 2016). For homologous parcels between left and right hemispheres to have the same hierarchical level in the heterogenous model, the T1w/T2w maps were symmetrized by averaging the T1w/T2w map values of the homologous parcels between left and right hemispheres.

### MEG Data Processing

We used resting-state MEG data of 89 subjects from the Human Connectome Project (HCP) (Larson-Prior et al., 2013; Van Essen et al., 2013). 3 runs of 6 minutes each were recorded per subject with a sampling frequency of 2034.5 Hz. Preprocessing included in the HCP release performed the following steps: removal of channels and segments as determined by the HCP quality assurance standards (Larson-Prior et al., 2013), bandpass (1.3–150 Hz) and notch (59–61 Hz, 119–121 Hz) filtering, and removal of non-brain components through independent component analysis. Moreover, a precomputed single shell volume conduction model was provided in this release.

Preprocessing followed the pipeline provided in the HCP dataset, the steps of which we summarize here. Source reconstruction was performed for approximately 8,000 vertices on the cortical surface, using the software provided by HCP and custom scripts written in Matlab and employing the Fieldtrip toolbox. Sensor data was bandpass filtered from 1.3 to 55 Hz and then projected to source space by synthetic aperture magnetometry (Vrba and Robinson, 2000). The sensor-covariance matrix was regularized by adding a value of 75% of its mean eigenvalue to the diagonal, and the noise covariance was assumed spherical. The direction of the source dipole was determined using a non-linear search in each dipole’s tangential plane to obtain the maximum signal-to-noise-ratio for the source power. After source reconstruction on the 8k-grid, source time courses were parcellated using the 360-area MSMAll atlas (Glasser et al., 2016). Each area’s time course was determined as the first principal component of its constituting voxels’ time courses.

Power spectra for each parcel were computed using Welch’s method with a frequency resolution of about 1 Hz and normalized such that the spectrum from 2–50 Hz sums to 1. For each parcel, they were then averaged across subjects and runs.

### Resting-State Network Assignments

We used network-level analyses to benchmark the hierarchical organization of empirical and simulated resting-state FC measure (i.e., the variation in sensory vs. association areas). Resting-state network (RSN) assignment was performed through community detection analysis on the correlation between resting-state fMRI time series of HCP dataset (Ito et al., 2017). The hierarchical categorization (i.e., sensory vs. association) of appropriate measures was calculated by averaging the values across 3 sensory networks (visual, auditory and somatomotor), and across 5 association networks (dorsal attention, ventral attention, default mode, fronto-parietal and cingulo-opercular).

### Large-Scale Computational Model

#### Synaptic dynamical equations

We adapted the biophysically-based large-scale computational model proposed by Deco et al. (2014b). This model reduces the complexity and the number of local microcircuit parameters in a spiking neural network model using a dynamical mean field approach (Wong and Wang, 2006). Exploiting the long time-constants of NMDA receptors, the local node model reduces from a large spiking neural network to a two-dimensional dynamical system.

Each cortical area is characterized by a microcircuit comprising a system of coupled excitatory and inhibitory populations. For each cortical microcircuit node *i* ∊ {1,…,*N*}, excitatory (*E*) and inhibitory (*I*) currents are given by:

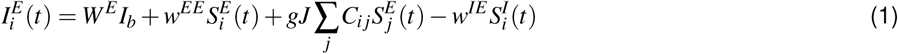

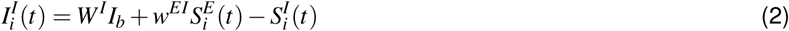

where *W^p^I_b_* and 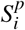, respectively, denote the background current and synaptic gating variable for each population *p* ∊ {*E*,*I*}, *g* is the global coupling parameter, *C_ij_* is the long-range structural connectivity strength from node *j* to node *i*, *J* is the effective NMDA conductance, *w^EE^* sets the local excitatory-to-excitatory strength, *w^EI^* sets the local excitatory-to-inhibitory strength, and *w^IE^* sets the local inhibitory-to-excitatory strength.

The firing rate of each population, *r^p^*, *p* ∊ {*E*,*I*} is computed using the transfer function 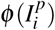:

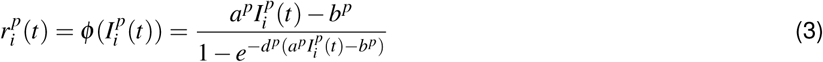

Finally, the synaptic gating variables obey:

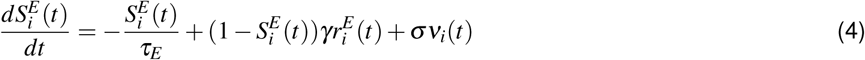

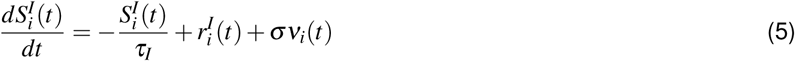

where *σ* is the standard deviation of the input noise taken from a random Gaussian process *v_i_* (Wong and Wang, 2006; Deco et al., 2014b).

Following Deco et al. (2014b), for each parameter set, the inhibitory-to-excitatory strengths *w^IE^* were adjusted to satisfy the condition 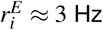. This was done by analytically solving for *w^IE^* to satisfy the self-consistency of Equations 1–5 at the steady-state condition with 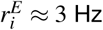, which corresponds to 〈*S^E^*〉 ≈ 0.164757 and 〈*I^E^*〉 ≈ 0.37738 nA:

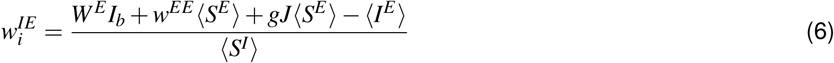

where the steady-state inhibitory synaptic gating variable 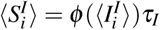 was estimated numerically by solving for 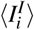:

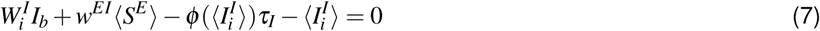

#### Hemodynamic equations

The synaptic activity of each cortical area is defined by the excitatory synaptic gating variable (*S^E^*). The excitatory synaptic activity of each area was transformed to a blood-oxygen-level dependent (BOLD) signal using the Balloon-Windkessel model (Friston et al., 2003), with parameter values from Obata et al. (2004). In the Balloon-Windkessel model, the hemodynamic response obeys using the system of equations:

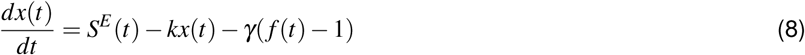

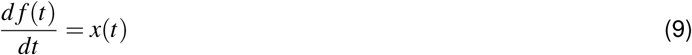

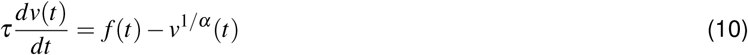

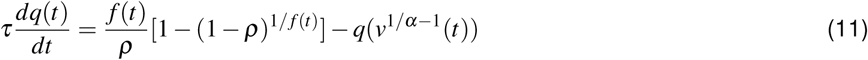

where *S^E^* is the excitatory synaptic gating variable, *x* is the vasodilatory signal, *f* is blood inflow, *v* is blood volume, and *q* is deoxyhemoglobin content. The parameters *ρ*, *V*_0_ and *τ* are the resting oxygen extraction fraction, resting blood volume extraction and hemodynamic transit time, respectively. The parameters *κ*, *gamma*, and *alpha* are rate of signal decay, rate of flow-dependent elimination and Grubb’s exponent, respectively. Based on the hemodynamic model, where *k*_1,2,3_ are dimensionless parameters, the BOLD signal is calculated as:

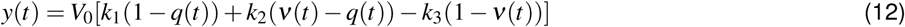

Values for all fixed parameters in Equations 1–12 are provided in **Table 1**.

**Table 1:** Fixed parameter values for synaptic and hemodynamics equations in the model.

| Synaptic model parameters |  |  |  |
| --- | --- | --- | --- |
| $I_b$ | 0.382 nA | | |
| $J$ | 0.15 nA | | |
| $\gamma$ | 0.641 | | |
| Excitatory populations |  | Inhibitory populations |  |
| $W^E$ | 1.0 | $W^I$ | 0.7 |
| $\tau^E$ | 0.1 s | $\tau^I$ | 0.01 s |
| $a^E$ | 310 nC<sup>-1</sup> | $a^I$ | 615 nC<sup>-1</sup> |
| $b^E$ | 125 Hz | $b^I$ | 177 Hz |
| $d^E$ | 0.16 s | $d^I$ | 0.087 s |
| Hemodynamic model parameters |  |  |  |
| $\rho$ | 0.34 | | |
| $\alpha$ | 0.32 | | |
| $V_0$ | 0.02 | | |
| $\gamma$ | 0.41 s<sup>-1</sup> | | |
| $\kappa$ | 0.65 s<sup>-1</sup> | | |
| $k_1$ | 7 $\rho$ | | |
| $k_2$ | 1.43 $\rho$ | | |
| $k_3$ | 0.43 | | |

#### Analytical Approximation of BOLD Functional Connectivity

The correlations between the excitatory populations of the cortical microcircuits can be approximated analytically by linearizing the system near a stable fixed point Deco et al. (2013; 2014b). In brief, the linearization of the model equations enables computationally efficient calculation of the synaptic FC matrix. However, in prior studies by Deco et al. (2013; 2014b), this calculation was not extended to fluctuations of the BOLD signal. We extended the linearization approximation to include the hemodynamic model for the BOLD signal.

The equations for synaptic dynamics (Equations 4–5) and the hemodynamic response (Equations 8–11) can be combined to form a single system 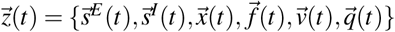 The details of the derivation for the moments of the synaptic equations were described previously in Deco et al. (2013; 2014b). Here, we derived the first moments of each hemodynamic quantity, which can be written as:

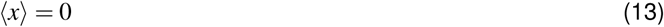

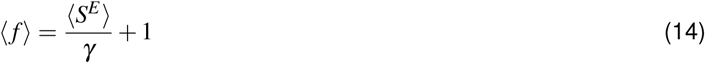

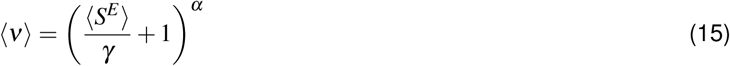

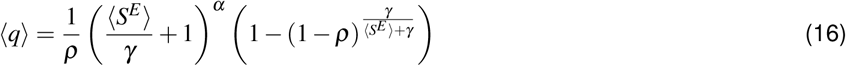

where 〈…〉 denotes the average over time.

Given the Jacobian of the extended system (**A**) and noise covariance matrix (**Q_n_**) at the stable fixed point, the system covariance matrix (**P**) can be approximated analytically by the Lyapunov equation:

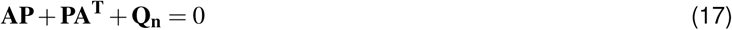

The Jacobian of the extended system involves the partial derivatives of the equations for synaptic dynamics (Equations 4–5) and the hemodynamic response (Equations 8–11). The Jacobian of the subset of the extended system describing synaptic dynamics (**A_syn_**) can be obtained from the partial derivatives of the system of equations (Equations 4–5) around the mean values:

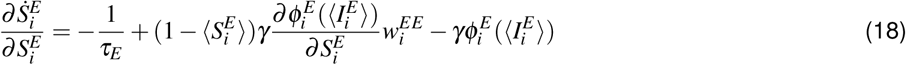

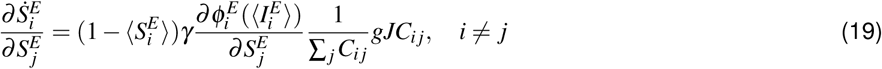

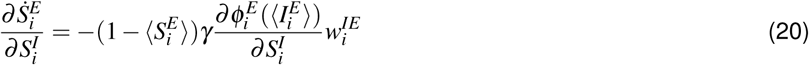

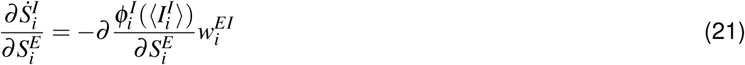

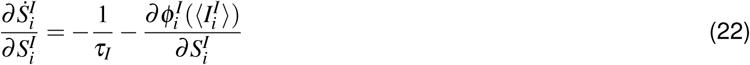

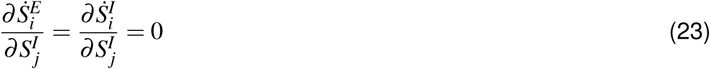

where 〈…〉 denotes the mean over time of each parameter. The Jacobian of the extended system (**A**) can be constructed by including the partial derivatives between the hemodynamic variables (Equations 8–11) and excitatory synaptic gating variables (Equations 4–5), which are given as:

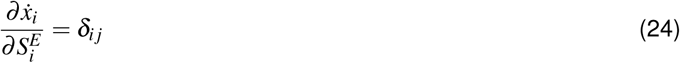

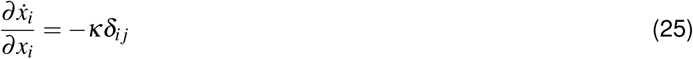

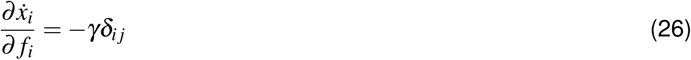

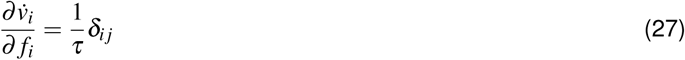

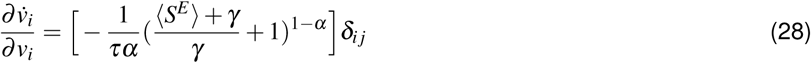

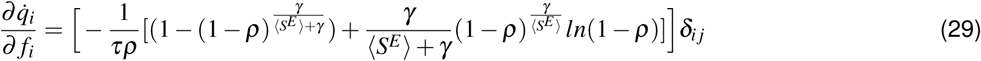

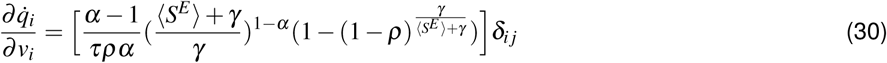

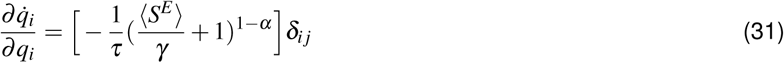

where *δ_ij_* = 0 if *i* ≠ *j*, and *δ_ij_* = 1 otherwise. The remaining partial derivatives are 0. Then, the covariance of the extended system **P** can be computed using the Lyapunov equation (Equation 17). Finally, the covariance of the BOLD signals is computed as:

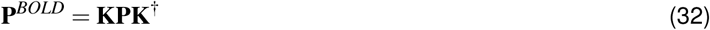

where **K** is the matrix comprising the terms describing the relationship between BOLD signal and hemodynamic quantities in Equation 12. The model-estimated BOLD FC between areas *i* and *j* can be estimated from the covariance matrix as:

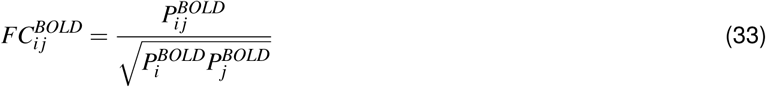

### Theoretical Characterization of the Model

To provide a mechanistic view of the local microcircuit dynamics, we performed theoretical analysis of the model from a dynamical systems approach. We first characterized the dynamical stability and the qualitative behavior of a single model node, with no external current (*I_ext_* = 0). For a set of local excitatory-to-excitatory and excitatory-to-inhibitory synaptic strengths (*w^EE^* and *w^EI^*, respectively), the feedback inhibitory-to-excitatory strength *w^IE^* is set to maintain an excitatory firing rate at *r^E^* ≈ 3 Hz (Deco et al., 2014b) (See section: Homogeneous and heterogeneous modeling paradigms). Therefore, although the phase diagram was constructed for *w^EE^* and *w^EI^*, the solution of the system implicitly depends on *w^IE^*.

The stability of the model was characterized by the largest eigenvalue *λ* of the Jacobian matrix **A_syn_**. We denote ℜ{*λ*} and ℑ{*λ*} as the real and imaginary parts, respectively, of the eigenvalue with largest real part. The fixed point dynamics are stable if ℜ{*λ*} < 0 and ℑ{*λ*} = 0, unstable if ℜ{*λ*} > 0 and ℑ{*λ*} = 0, stable spiral if ℜ{*λ*} < 0 and ℑ{*λ*} ≠ 0, and unstable spiral if ℜ{*λ*} > 0 and ℑ{*λ*} ≠ 0. The system exhibits pitchfork bifurcation when ℑ{*λ*} = 0 and a Hopf bifurcation when ℑ{*λ*} ≠ 0.

For *w^EI^* values between 0.001 and 5, we numerically solved the equations for *w^EE^* when ℜ{*λ*} = 0 and ℑ{*λ*} = 0 for *w^EE^* to determine critical points. The natural frequency of the system was calculated as |ℑ{*λ*}|/2*π*.

To illustrate optimal model parameters projected onto the phase plane, we adjusted the external input according to the total synaptic input to each area such that *I_ext_* = *gJ*〈*S^E^*〉 where *g* is the global coupling parameter and *J* is the long-range NMDA conductance, and 〈*S^E^*〉 is the average synaptic gating variable.

### Homogeneous and Heterogeneous Modeling Paradigms

In both modeling paradigms, all parameters were kept constant except local excitatory-to-excitatory synaptic strength *w^EE^*, excitatory-to-inhibitory synaptic strength *w^EI^*, and global coupling parameter *g*. In the homogeneous model, the values of recurrent excitatory synaptic strength *w^EE^* and excitatory-to-inhibitory synaptic strength *w^EI^* were assigned globally; i.e., same value for each area. In the heterogeneous model, *w^EE^* and *w^EI^* were defined by a map based on the hierarchical level of each area.

We showed that the T1w/T2w map is characterized by high values in sensory regions and low values in association regions (**Figure 1**). In the heterogeneous model, we introduced hierarchical heterogeneity using the median T1w/T2w map values computed across 334 subjects. The raw T1w/T2w map values across brain areas exhibit a positively skewed distribution (Glasser et al., 2014). For model parameterization, we transformed the raw T1w/T2w map values into a hierarchical heterogeneity map values which are more uniformly distributed, rescaled between 0 and 1, and inverted such that high- (low-) T1w/T2w areas have low (high) hierarchical heterogeneity map values (**Figure S2**). Specifically, hierarchical heterogeneity map value *h_i_*, for each *i*, was determined by:

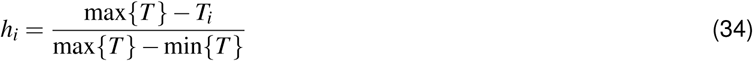

*T_i_* is calculated from the raw T1w/T2w value 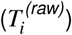 using the error function erf: 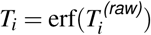. Local microcircuit parameters *w^EE^* and *w^EI^* were linearly scaled by *h_i_* map values:

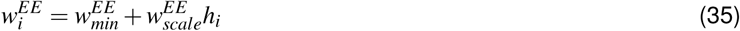

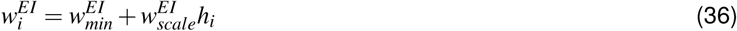

where the subscript *min* denotes the minimum parameter value and the subscript *scale* denotes the scaling factor that defines the steepness of the hierarchical heterogeneity map.

We chose to use white-matter/gray-matter seeding for diffusion tractography, as this improves agreement of within-hemisphere connectivity with tracer-measured connection strengths (Donahue et al., 2016). We found that the correlation between SC and FC is primarily driven by intra-hemispheric connections (**Figure S1**). Therefore, we focused on modeling and analyzing the network structure of FC within each hemisphere due to intra-hemispheric interactions. In addition, on our high-performance computing cluster, the runtime of a single execution of the model was approximately 1 second for the intra-hemispheric model (180 regions), whereas it exceeded 10 seconds for the entire cortex (360 regions). For these reasons, we excluded inter-hemispheric connections and simulated the left and right hemisphere models separately. In both homogeneous and heterogeneous paradigms, the models comprised two intra-hemispheric compartments in which the local microcircuit parameters in homologous areas are identical but the two hemispheres do not interact.

### Optimization of Model Parameters

We estimated the optimal model parameters for homogeneous and heterogeneous models. The homogeneous model included four free parameters: *w^EE^*, *w^EI^*, and *g_l_*, and *g_r_*. The heterogeneous model parameters included six free parameters: 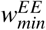, 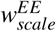, 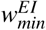, 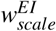 and *>g_l_*, and *g_r_*.

Bayesian statistics has substantial advantages over point estimates of model parameters because it provides a better description by estimating the full posterior probability distributions of the model parameters. However, estimation of the model FC is based on a stochastic dynamical system. Therefore, it is not possible to analytically solve for the likelihood function. For this reason, we used Approximate Bayesian Computation (ABC), which approximates the likelihood function by minimizing a distance measure, to estimate the optimal model parameters. To find the model parameters that minimize the distance between empirical FC (*FC*) and model 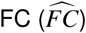, we used adaptive hierarchical Population Monte Carlo (hPMC) (Beaumont et al., 2009; Turner and Van Zandt, 2014). Since the variations across subjects are removed in average FC, we defined the distance measure based on individual FCs. Specifically, we estimated parameters which minimized the average empirical and model FC distance across the entire population (Turner and Van Zandt, 2014) (**Figure S2**).

In the initial step, each parameter value was drawn from its prior distribution (**Table 2**). In subsequent steps, the parameters were drawn from a proposal distribution. The prior distributions were informed by the phase-diagram (**Figure 7** and **Figure S5**).

**Table 2:** Prior distributions for hPMC model fitting. U(MIN, MAX) denotes a uniform probability distribution between MIN and MAX.

| Parameter ($\theta$) | Homogeneous | Heterogeneous |
| --- | --- | --- |
| $w_{min}^{EI}$ | U(0.001, 5.0) | U(0.001, 2.0) |
| $w_{min}^{EE}$ | U(0.001, 15.0) | U(0.001, 5.0) |
| $g_l, g_r$ | U(0.001, 5.0) | U(0.001, 2.0) |
| $w_{scale}^{EI}$ | | U(0, 2.5) |
| $w_{scale}^{EE}$ | | U(0, 15.0) |

In the PMC approach, a particle is defined as a set of model parameters that is drawn from the proposal distribution. For each particle, the model FC 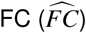 was calculated according to Equation 26, if the system of equations is dynamically stable. Then, we calculated the distance (*δ*) as:

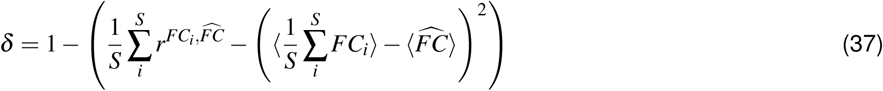

where *r* is the Pearson correlation coefficient, *S* ∊ { 1, …, 334} is the number of subjects, and 〈〉 denotes the average FC across regions. The first term in the parentheses of Equation 37 quantifies the average Pearson distance between the model FC and subject FCs. The second term is an additional cost term quantifying the difference between mean FCs (since Pearson correlations involves mean-subtraction), ensuring that the mean model FCs do not diverge from the mean empirical FCs.

The particles were rejected if the distance, *δ*, was larger than the rejection threshold *ε*. Initially, the rejection threshold was defined as the Pearson correlation distance between the empirical FC and SC. In subsequent iterations, the rejection threshold was iteratively adjusted according to the first quantile of accepted distances in the previous iteration (Drovandi and Pettitt, 2013). The algorithm was run until the number of accepted particles exceeded the minimum sample size (*N*). For the rest of the iterations (*T*), samples were drawn from the proposal distribution:

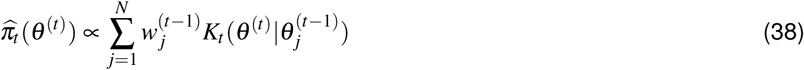

where the importance weight 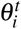 for an accepted particle 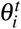 is

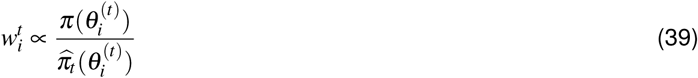

and the random walk kernel (*K_t_*) is defined as:

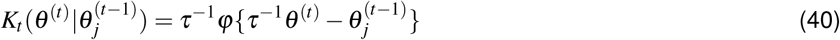

Where *φ* is standardized (multivariate) normal density and *τ* is the scaling factor, which is calculated as twice the weighted empirical covariance: *τ*^2^ = 2Cov(*θ*).

In both homogeneous and heterogeneous models, the minimum sample size was set to *N* = 1000 and the maximum number of iterations was set to *T* = 100. The algorithm was terminated if it reached either the maximum number of iterations or the acceptance rate (i.e., *N*/*N_total_*_)_ was lower than 0.001, indicating convergence. The homogeneous model converged after 75 iterations and the heterogeneous model converged after 76 iterations due to low acceptance rate, but the distance between empirical and model FC stabilized after approximately 50 iterations (**Figure S3**).

To test that the improved fitting in the heterogeneous model is not due to overfitting with the higher number of degrees of freedom in the model, we performed leave-p-out cross-validation. We repeated the optimization procedure for both homogeneous and heterogeneous models (using left cortical hemisphere, minimum sample size *N* = 200, maximum number of iterations *T* = 50) on 267 of 334 subjects, after holding out 20% of the subjects randomly. Then, we calculated the model fit on the held-out test subjects.

In addition to leave-p-out cross-validation, we compared the T1w/T2w-derived hierarchical heterogeneity map against surrogate maps with spatial autocorrelation structure matched to the hierarchical heterogeneity map, as described in the following section (**Figure 3**). We repeated the procedure for 500 surrogate heterogeneity maps. For computational efficiency, we ran the optimization for left cortical hemisphere, with a minimum of 500 particles for at most 50 iterations.

We also performed this fitting procedure for procedure for T1w/T2w map derived hierarchical heterogeneity map under the same conditions. The best fit for re-optimized T1w/T2w-derived hierarchical heterogeneity map was same as that for the main optimization of the heterogeneous model (i.e., left-right concatenated). For all maps, we ensured that the similarity between empirical and model FC stabilized within 50 maximum iterations (**Figure S3**).

### Surrogate Heterogeneity Map Generation

First we characterized the spatial autocorrelation structure of median empirical cortical T1w/T2w map using a spatial lag model. We fit the data using a spatial lag model of the form **y** = *ρ***Wy**, where **y** is a vector of first Box-Cox transformed and then mean-subtracted map values. The Box-Cox transformation was first applied to the maps so their values were approximately normally distributed. **W** is the row-normalized weight matrix with zero diagonal and off-diagonal elements proportional to 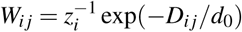, where *D_ij_* is the surface-based geodesic distance between cortical areas *i* and *j*, and *z_i_* ≡ ∑*_j_* exp(−*D_ij_*/*d*_0_) is a row-wise normalization factor. Weights *W_ij_* define the fraction of spatial influence on area *i* attributable to area *j*.

Two free parameters *ρ* and *d*_0_ are estimated by minimizing the residual sum-of-squares (Anselin, 2001). Using best-fit parameter values 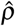 and 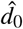, surrogate maps **_y_**_surr_ are generated according to 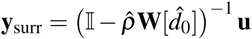, where 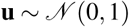. For the Box-Cox normalized T1w/T2w values, this fit yielded *d*_0_ = 6.152 mm, and *ρ* = 1.050. To match surrogate map values distributions to the distribution of values in the corresponding empirical map (e.g., the T1w/T2w map), rank-ordered surrogate map values were re-assigned the corresponding rank-ordered values in the empirical map. Note that this approach to surrogate data generation approximates a spatial autocorrelation-preserving permutation test of the empirical neuroimaging map.

Using these surrogate maps we construct null distributions for *N* = 500 models and report significance values as the proportion of samples in the null distributions whose model fit value is greater than or equal to that obtain from the model using the T1w/T2w-derived hierarchical heterogeneity map. where *d_ij_* is the average surface-based geodesic distance between the grayordinates of cortical parcels *i* and *j*. To calculate geodesic distances, we used the 32k-vertex midthickness surface mesh in the HCP atlas.

### Spatial Auto-Regressive Modeling

Significance values indicated by the number of stars reported on bar plots for MEG band-limited power (BLP) maps and the T1w/T2w map correlations were corrected to account for spatial autocorrelation structure in parcellated T1w/T2w maps and BLP maps. Because physical quantities must vary smoothly and continuously in space, measurements recorded from proximal cortical areas tend to be more similar than measurements recorded from distal areas of cortex. This departure from the assumption of independent observations biases calculations of statistical significance. To model this spatial autocorrelation, we used a spatial lag model (SLM) commonly applied in the spatial econometrics literature (Anselin, 2001), of the form *y* = *ρWy* + *Xβ* + *ν*, where *W* is a user-defined weight matrix implicitly specifying the form of spatial structure in the data, and *ν* is normally distributed.

To implement a spatial lag model in the python programming language, we used the maximum likelihood estimation routine defined in the Python Spatial Analysis Library (*pysal*) (Fischer and Getis, 2010). We first determined the surface-based spatial separation between each pair of cortical parcels by computing the mean of the pairwise geodesic distances between each vertex in parcel *i* and each vertex in parcel *j*, from which we constructed a pairwise parcel distance matrix, *D*.

We estimated the spatial autoregression parameters 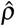 and 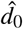 for each empirical MEG BLP map. Then, using the OLS estimate of the spatial autocorrelation scale from the fit to the empirical MEG BLP maps, we calculated the elements of the spatial weight matrix, *W_ij_* = exp(−*D_ij_*/*d*_0_). Finally, we fit the SLM to empirical MEG BLP maps, using the maximum likelihood estimator routine (`pysal.spreg.ml_lag.ML_Lag`) in *pysal*. In Figure 8, p-values indicated by the number of stars correspond to p-values for model parameter *β* defined above.

Using the OLS estimate of the spatial autocorrelation scale from the fit to the empirical BLP, we calculated the elements of the spatial weight matrix, *W_ij_* = exp(−*D_ij_*/*d*_0_). Finally, we fit the SLM to BLP values, using the maximum likelihood estimator routine (`pysal.spreg.ml_lag.ML_Lag`) in *pysal*. P-values indicated by the number of stars in the bar plots of T1w/T2w map correlations correspond to p-values for model parameter *β* defined above.

Of note, spatial autoregressive model parameters do not have the same interpretation as they do in OLS regression. The parameter *β* reflects the direct (i.e., local) impact on the dependent variable *y* due to a unit change in the independent variable *x*. In addition, because of the underlying spatial structure, the direct impact of *x_i_* on *y_i_* results in an indirect effect of *y_i_* on neighboring *y_j_*. Therefore *β* cannot be interpreted as a corrected, global correlation coefficient, and we restrict our use of the SLM to correcting for the biasing effect of spatially autocorrelated samples on reported significance values. Finally, the significance values were corrected for multiple comparisons using the false discovery rate (FDR).

### Examination of Confounding Variables

We used regression analysis to test whether the improved rs-FC fit of the heterogeneous model, relative to the homogenous model, may be driven by fitting of FC components attributable to non-neural confounding factors such as head motion, heart rate variations, and respiration (Power et al., 2017). The movement parameters and physiological logs were provided by the HCP dataset. The head motion of each subject was calculated as the mean absolute movement and mean relative movement. Heart rate and respiration variations were calculated as the total variance of each variable across sessions. To test whether the relationship between movement parameters and model fit can explain the improved performance of the heterogeneous model, we constructed a linear regression model (*Y^(full)^*):

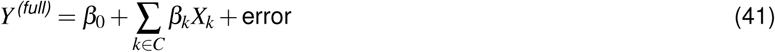

Each subject yields a data point in *Y*, defined as the difference in model fit for the heterogeneous model compared to the homogeneous model (i.e., *r*_heterogeneous_ − *r*_homogeneous_). *X_k_* denotes the z-score of four subject-specific measures non-neural confounding factor *k* in the set of four denoted *C*: mean absolute head movement, mean relative head movement, heart rate variance, and respiration variance.

We compared the full regression model *Y^(full)^* above to a reduced model that corresponds to the difference in model fit being independent of confounds: *Y^(reduced)^* = *β*_0_ + error. Through comparison of the full and reduced regression models, we examined contribution of the non-neuronal confounding factors on the improved model fit in the heterogeneous model. We compared the fraction of variance in the data which is captured by the model, calculating the ratio of explained variance for the full vs. reduced regression models, which we found to be very small.

### Within-Network and Across-Network Fitting

To study the differences between the performances of homogeneous and heterogeneous models, we quantified the model fitting for the FCs within and across 8 canonical regarding the resting state networks (RSNs). The within-network fit for network *N* was calculated as:

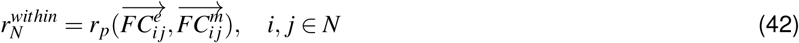

where 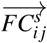, *s* ∈ {*e* : empirical, *m*: model} is vector comprising the mean functional connectivity values between regions *i* and *j*, and *r_p_*(*x*, *y*) denotes the Pearson correlation coefficient between variables *x* and *y*. Similarly, the across-network fit for network *N* is defined as:

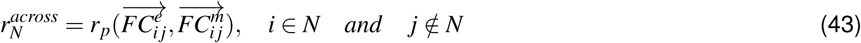

### Global Brain Connectivity

We tested whether hierarchical heterogeneity can also explain rs-FC measures which exhibit a hierarchical preference. One such measure is the Global Brain Connectivity (GBC) (Cole et al., 2010). Here, we adapted GBC (Cole et al., 2010), to parcellated rs-FC matrices. The GBC of region *i* was defined as the average FC between region *i* and the rest of the regions:

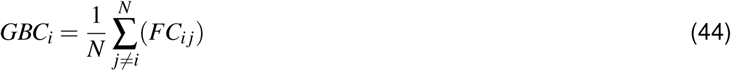

where *N* is the number of brain region. GBC is equivalent to the graph-theoretic measure of node strength normalized by total number of nodes.

### Inter-Individual Dissimilarity

To characterize variation in rs-FC patterns of each region across subjects, we adapted inter individual variability as proposed in Mueller et al. (2013). Since this measure reflects the dissimilarity across rs-FC patterns, we refer to it as a measure of dissimilarity instead of variability. In empirical data, we refer to the measure as inter-individual dissimilarity, whereas in the model, we refer to it as inter-particle dissimilarity. The inter-individual/particle dissimilarity *V_i_* is calculated as:

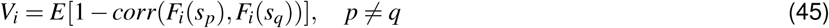

where *F_i_*(*s*) is a vector of rs-FC balues between region *i* and rest of the regions in subject/particle *s*, *corr*(*x*,*y*) is the Spearman rank-correlation between variables *x* and *y*, and *E*[.] denotes expected value. Due to the analytical approximation of the model FCs, the scales of inter-individual and inter-particle dissimilarity were different. To normalize the scales of both measures, we z-scored the final spatial maps of inter-individual/particle dissimilarity. Of note, MMSAll registration does not produce perfect cross-subject alignment due to atypical topologies and therefore some portion of inter-individual dissimilarity is due to misregistration of areas to the atlas (Bijsterbosch et al., 2018).

### Spectral Characterization of the Model

As described in Deco et al. (2014b), the power spectrum of the model can be analytically approximated around its fixed points. The cross-spectrum Φ as a function of frequency *ω* can be calculated by:

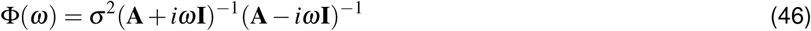

where **A_syn_** is the Jacobian of the system of synaptic variables described in equations 1-5, *σ* is the standard deviation of the input noise, *i* is 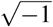, and **I** is identity matrix.

The power spectral density (PSD) was computed as the diagonal elements of Φ(*ω*) for frequencies *ω* ∊ (0.01 … 75 Hz). The PSDs of left and right hemisphere models were normalized by dividing by total PSD. The PSDs were reported in dB (10log_10_). The band-limited power (BLP) was calculated by averaging the PSDs within 5 frequency bands, which were defined as slow in 0.001–2 Hz (only in model), delta (*δ*) in 2–4 Hz, theta (*θ*) in 4–8 Hz, alpha (*α*) in 8–15 Hz, beta (*β*) in 15–35 Hz, and gamma (*γ*) in 35–50 Hz. The similarities between empirical and model BLPs, and empirical BLPs and the T1w/T2w-derived hierarchical heterogeneity map were quantify using Spearman rank correlation coefficient.

To study the role of long-range connections on the behavior of the system, we estimated the PSDs of the model without long-range connections (**Figure S6**). Briefly, we replaced the term 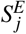 in Equation 1 with the average excitatory synaptic gating variable 〈*S^E^*〉, such that the long-range connection term reduced to an external synaptic input, i.e., *I_ext_* = *gJ*〈*S^E^*〉.

### Numerical Simulations

To show the validity of analytical approximation of BOLD FC and PSD, we employed full simulations of the model. We integrated synaptic Equations 1–5 and BOLD Equations 15–19 using Euler method with *dt* = 0.1 ms. The input noise levels were defined as *σ* = 10^−5^. To calculate BOLD FC, we simulated the BOLD signals for 870 seconds to match experimental recording sessions. The first 6 seconds of simulations were discarded to ensure that the system’s state converges to the neighborhood of the baseline fixed point. The simulated BOLD signals were downsampled using 0.72-s resolution resulting in 1200 TRs as in the empirical BOLD data from the HCP. To calculate simulated PSDs, we simulated the excitatory synaptic gating variables for 170 seconds, and later downsampled to 10 ms after removing the first 6 seconds of simulations. In total, 10 simulations were performed for 20 random iterations using particles that were drawn from the posterior distribution.

The same simulation protocol was repeated using synaptic delays. The distance between brain regions was defined by the average pairwise euclidean distance between two parcels. The conductance velocity was defined as *v* = 6*m*/*s*, which is consistent with empirically observed values (Horowitz et al., 2015) and with previous modeling studies (Cabral et al., 2011).

### Statistical Tests

As a statistical test for the difference between similarities of the measures derived from homogeneous and heterogeneous models to the empirical data, we employed dependent correlations test. Dependent correlations test (DepCor) provides a non-parametric test to compare the correlations of two variables against a common dependent variable based on bootstrapping (Wilcox, 2016). The DepCor tests were employed directly when Spearman rank correlation was used a measure of similarity (i.e., GBC and BLP). When Pearson correlation was used as a measure of similarity (i.e., FC, within-/across-network FC, inter-individual/particle dissimilarity), skipped Pearson correlation was used to perform DepCor tests due to sensitivity of Pearson correlation to possible outliers during bootstrapping. In the main results, only Pearson correlations were reported and using Pearson correlation instead of skipped Pearson correlation did not change the results.

To test for differences between the average measures in sensory and association areas in empirical data (i.e., T1w/T2w maps, GBC, inter-individual dissimilarity), we used the Wilcoxon signed-rank test. The Wilcoxon signed-rank test provides a non-parametric measure for the difference between the ranks of repeated measurements of two related samples.

The surrogate heterogeneity maps were used to generate various sample null distributions for statistics (e.g., model FC fitting). Reported significance values were calculated as the proportion of samples in the null distributions whose values exceeded that of the test statistic.

### Data and Software Availability

Custom modeling and analysis codes written in Python will be freely available upon acceptance. All results derive from the freely available HCP dataset. Parcellated maps and connectivity matrices related to this study will be freely available via the BALSA database (https://balsa.wustl.edu/) upon acceptance.

---

**Figure S1:**
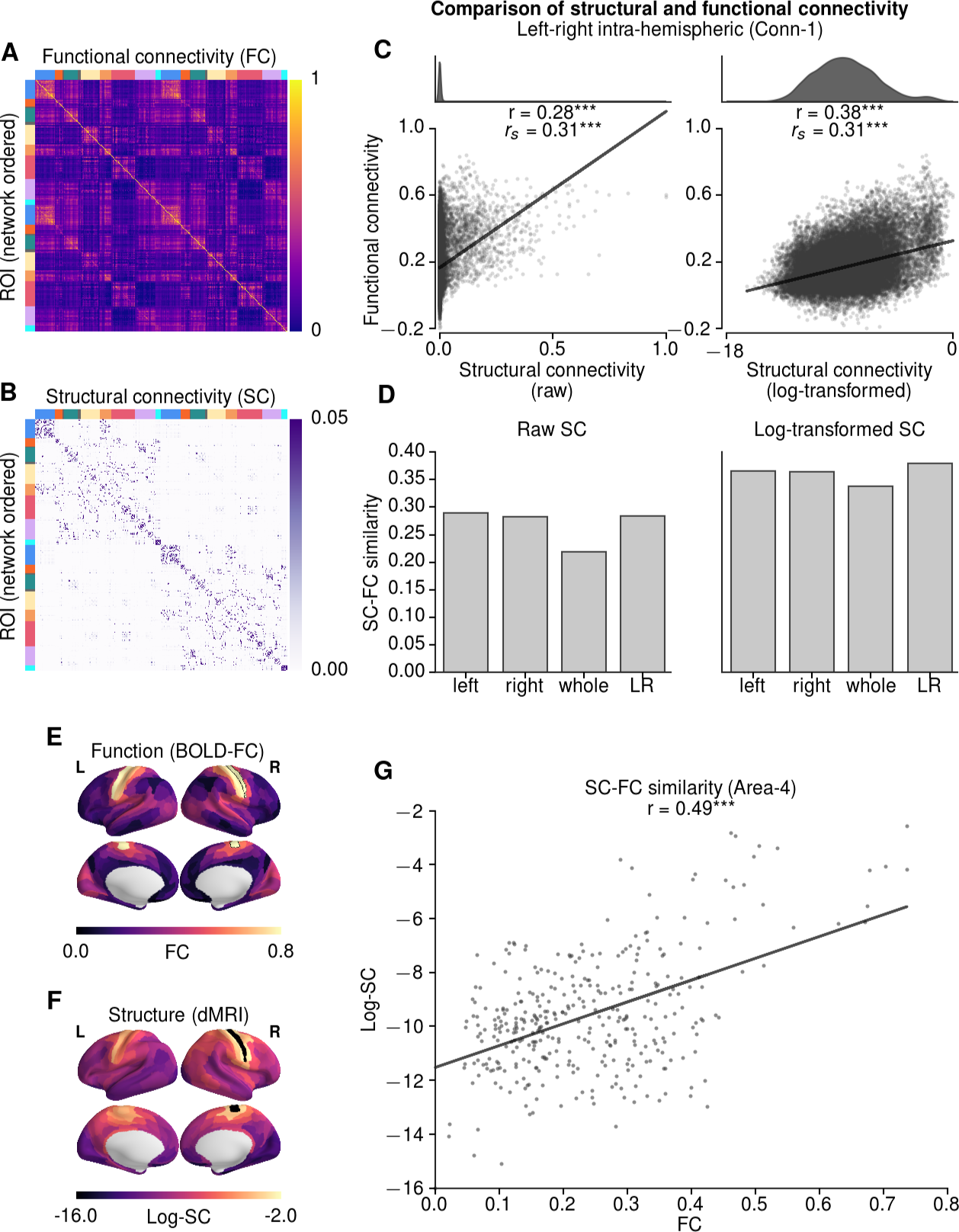
Correspondence Between Empirical Structural and Functional Connectivity. **(A,B)** Group-averaged bilateral structural connectivity (SC) derived from dMRI and functional connectivity (FC) derived from resting-state fMRI. The matrices are ordered by resting-state networks within each hemisphere (colored as in Figure 1). **(C)** Correlation between average FC and SC for intra-hemispheric connections, for raw SC weights (left) and log-transformed SC weights (right). Consistent with previous literature, we found a significant relationship between dMRI-derived SC and rs-FC (*r_s_* = 0.31, *p* < 10^−5^). **(D)** Correlation between group-averages FC and SC for intra-hemispheric (left, right), whole-brain (i.e., including intra- and inter-hemispheric connections), and left-right concatenated matrices. The SC-FC correlation is lower for the whole-brain matrix which includes inter-hemispheric connections. **(E,F)** FC and SC topographies with area 4 in the right hemisphere as an example seed seed region. **(G)** Correlation between whole-brain FC and log-transformed SC for the example seed region, right-hemisphere area 4 (*r* = 0.49, *p* < 10^−5^). *** indicates *p* < 0.0001.

**Figure S2:**
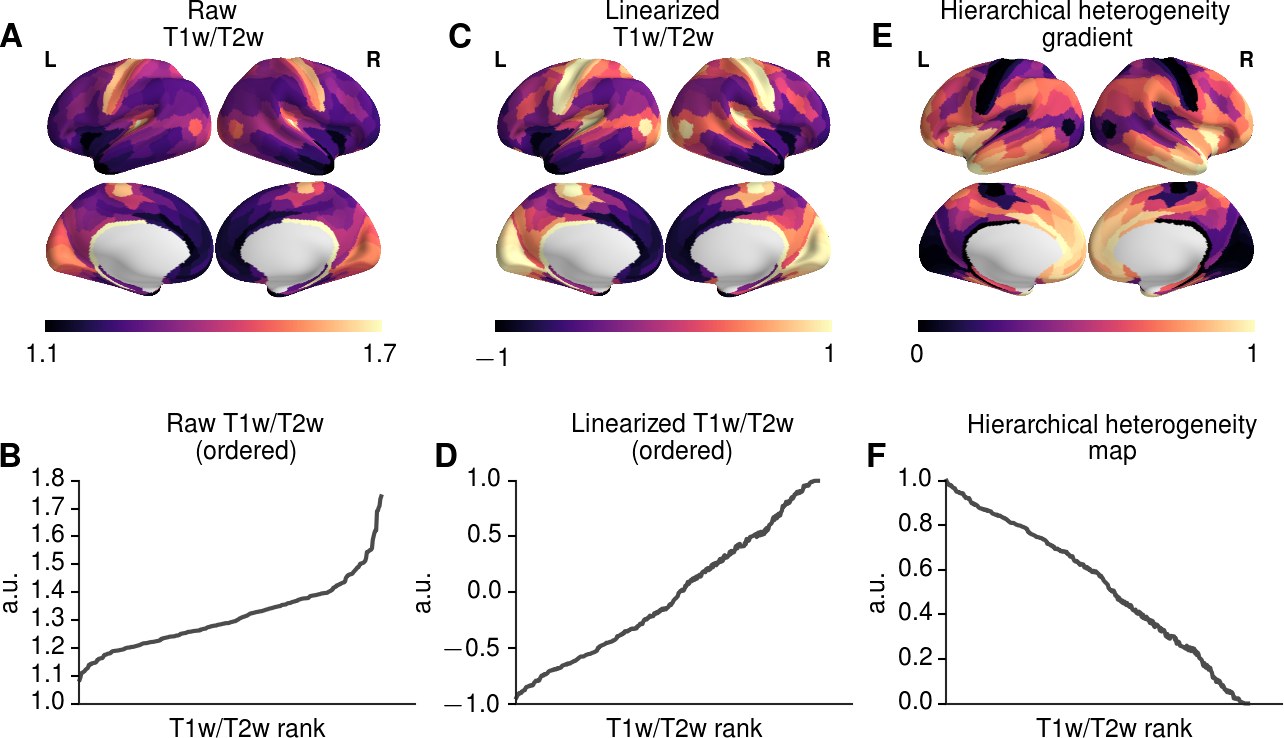
Calculation of the T1w/T2w-Derived Hierarchical Heterogeneity Map. **(A,B)** Rank ordering of T1w/T2w map values. **(C,D)** Rank ordering of linearized T1w/T2w map. Linearization is performed by transforming values with the error function. **(E,F)** Rank ordering of hierarchical heterogeneity map values used in the model. After linearizing T1w/T2w map values (*T_i_*), were normalized and inverted between 0 and 1 according to: 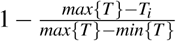.

**Figure S3:**
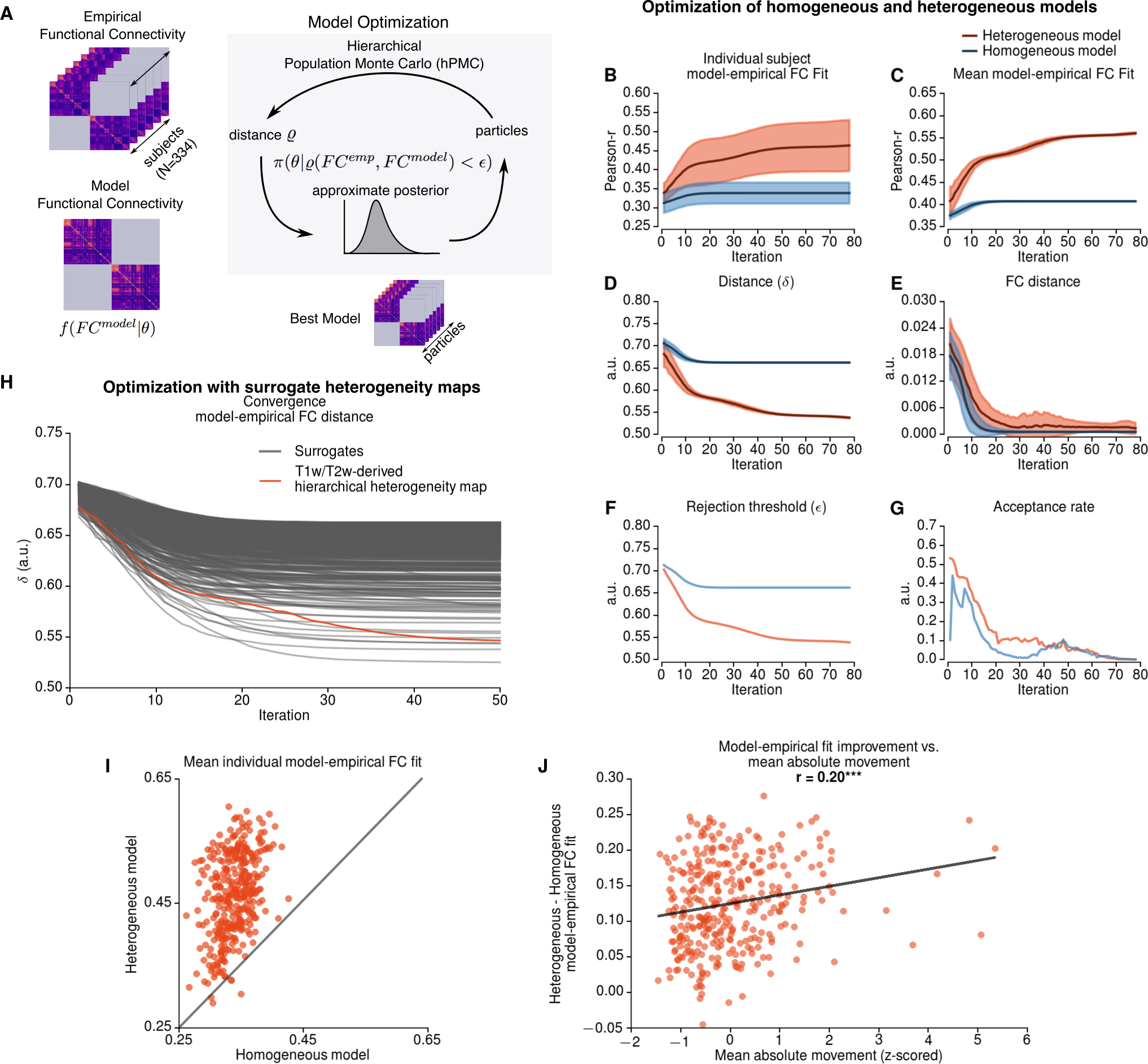
Optimization of Model Parameters. **(A)** Schematic for Approximate Bayesian Computation (ABC) via hierarchical Population Monte Carlo (hPMC). After sampling particles (a set of model parameters *θ*) from the proposal distribution and estimating the model FC, we calculated the average correlation (*r*) between subject FCs and the model FC. The distance measure *δ* was defined as 1 *–* (*r*+*c*), where *c* is an additional cost which controls for the mismatch between the model and empirical FC. A particle was accepted if the distance was smaller than the threshold (*ε*), which was decreased at each iteration. The particles were sampled until 1000 particles satisfied the threshold. Accepted particles were used to update the proposal distribution for the next iteration. The convergence criterion for terminating the optimization was an acceptance rate being lower than 0.001. **(B–G)** The evolution of optimization parameters across iterations for the homogeneous (blue) and heterogeneous (red) models. **(B)** Average correlation between model FC and individual subject FCs. **(C)** Average correlation between model FC and group-averaged empirical FC. **(D)** The distance measure *δ*. **(E)** The distance between the average model FC and average FC. **(F)** Rejection threshold. **(G)** Acceptance rate. The model fits were stabilized after 20 iterations for homogeneous model and 50 iterations for heterogenous model (**B–E**). The acceptance rate falls below 0.001 after 70 iterations for both models. **(H)** The evolution of distance between empirical and model FCs for the T1w/T2w map-derived heterogeneity map (red) and surrogate heterogeneity maps (gray). For all surrogate maps, the similarity between model and empirical FC stabilizes within 50 iterations. Shaded regions indicate standard deviations across particles. **(I)** Improved individual subject model fit in heterogeneous model compared to homogeneous model. For the majority of the subjects (329 of 334) the heterogeneous model performed better than homogeneous model. **(J)** The relationship between model improvement and mean absolute movement across subjects exhibited a weak but statistically significant correlation.

**Figure S4:**
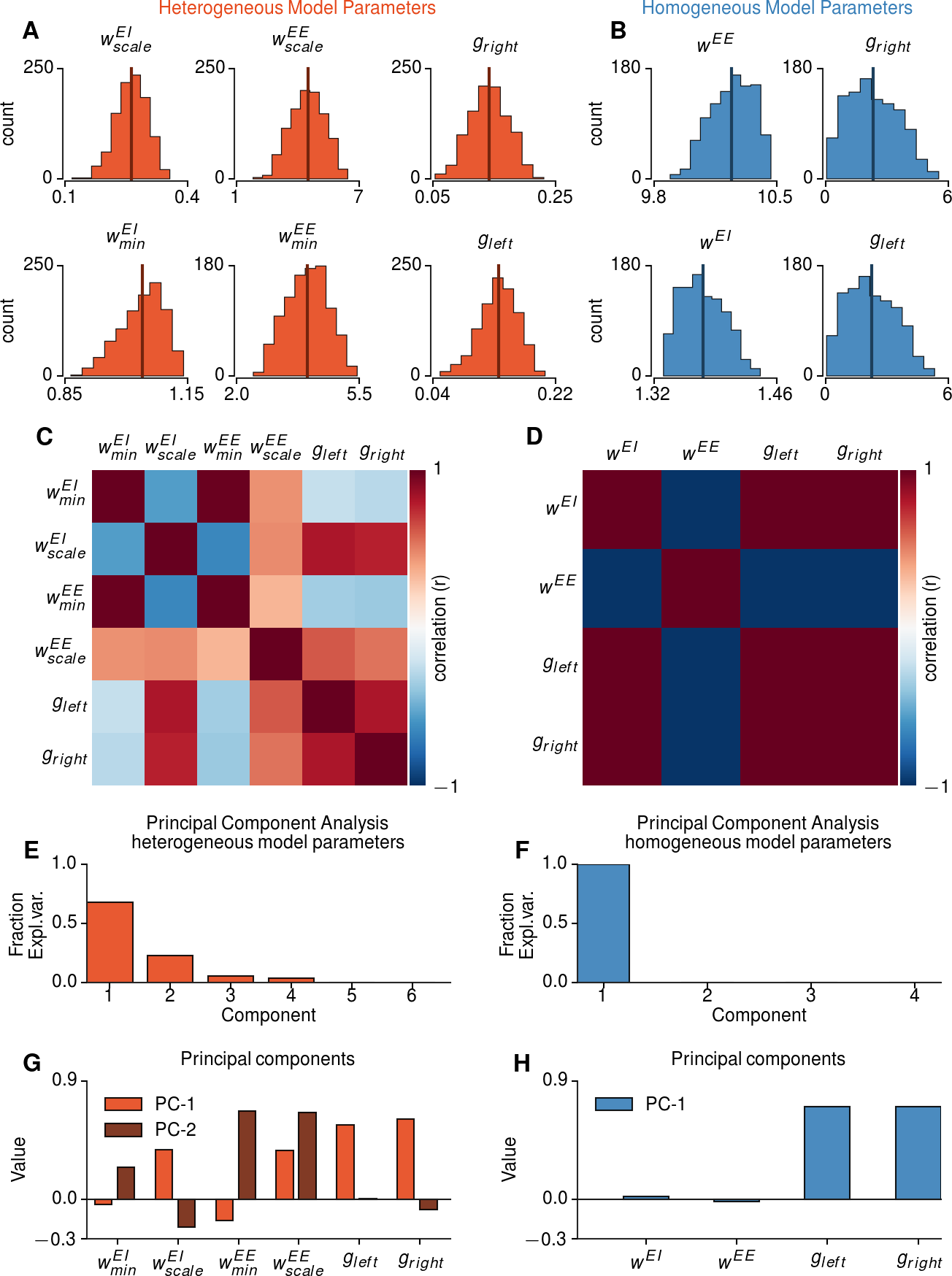
Relationships Between Model Parameters. **(A)** Marginal posterior distributions for heterogeneous model parameters. *w^EI^* intercept was 1.038 ± 0.053 with a scaling factor of 0.271 ± 0.041, and *w^EE^* intercept was 3.994 ± 0.59 with a scaling factor of 4.653 ± 0.861. The global coupling parameters were 0.143 ± 0.030 for left hemisphere and 0.148 f 0.034 for right hemisphere. Values are mean f std. dev. **(B)** Marginal posterior distributions for homogeneous model parameters. *w^EE^* = 10.168 ± 0.12 and *w^EI^* = 1.39 ± 0.023. The global coupling parameters were 2.956 ± 1.125 for left hemisphere and 3.064 ± 1.168 for right hemisphere. **(C)** The correlations between marginal posterior distributions for heterogeneous model parameters. There are strong correlations between the parameter 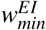, 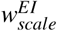, 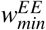. Similarly, the global coupling parameters of left and right hemispheres, *g_left_* and *g_right_*, are strongly correlated. The scaling factor 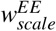 did not show strong correlations with parameters other than *w^EI^*. This suggests that the scaling of recurrent excitatory strengths along the hierarchy axis has a unique contribution to the variability across particles. **(D)** Correlations between marginal posterior distributions for homogeneous model parameters. Homogeneous model parameters are very strongly correlated with each other. **(E-H)** Principal component analysis (PCA) of the optimal model parameters. PCA was applied to the distribution of particles drawn from the posterior, with their parameter values normalized by the population mean for each parameter. The fraction of explained variance by the top principal components, for heterogeneous model **(E)** and homogeneous model **(F)**. The variation in heterogeneous model is explained by 4 components **(G)**. 100% of the variation in homogeneous model parameters is explained by a single dimension **(H)**.

**Figure S5:**
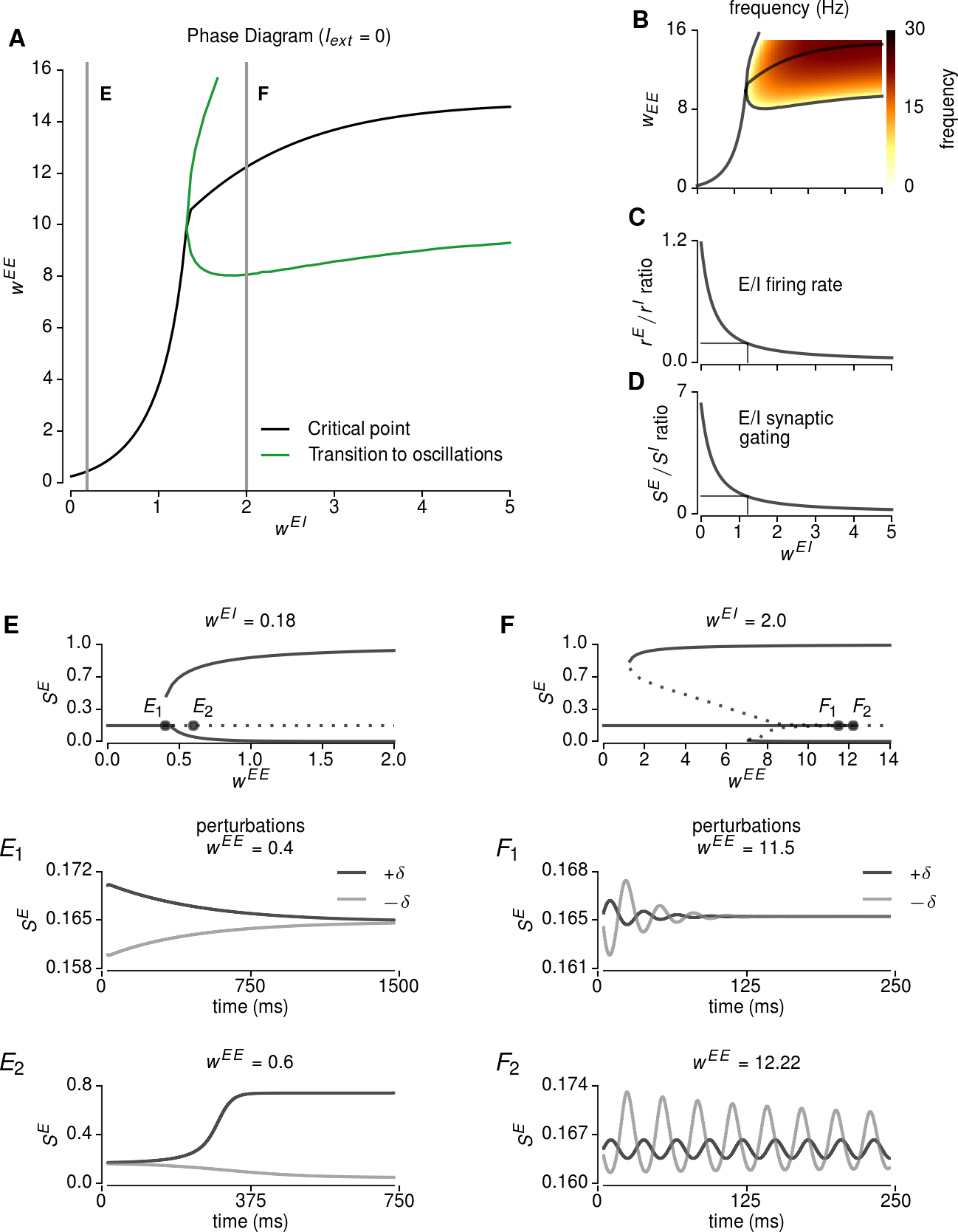
Dynamical Systems Analysis of a Local Excitatory-Inhibitory Node. **(A)** The extended phase diagram for *w^EE^* and *w^EI^*. For low values of *w^EI^* the model exhibits a single pitchfork bifurcation, whereas for high values the system exhibits oscillatory activity. **(B)** Intrinsic frequencies of the system calculated from imaginary parts of the eigenvalues of the Jacobian matrix. For high values of *w^EI^* and *w^EE^*, the system generates oscillations with intrinsic frequencies between 0 and 30 Hz. **(C-D)** The ratio of excitatory to inhibitory firing rates (**C**) and synaptic gating variables (**D**) as a function of excitatory-to-inhibitory strength *w^EI^*. For both models, the optimal parameter range was near the critical point at which the system exhibits a oscillatory activity. In this regime, the ratio between excitatory and inhibitory synaptic gating variables is approximately equal to 1 (i.e., the excitatory and inhibitory synaptic activities are balanced), while the firing rate of inhibitory neurons is higher than excitatory neurons, consistent with cortical physiological recordings. **(E)** The bifurcation diagram for *w^EI^* = 0.18 (i.e. the value proposed in Deco et al. (2014b)). The perturbations (±*δ*) around fixed point for *w^EE^* = 0.4 *E*_1_ and *w^EE^* = 0.4 *E*_2_. Before bifurcation, the synaptic gating variable returns to its steady state value *E*_1_. After bifurcation, the synaptic gating variable moves towards up- and down-attractor states *E*1. **(F)** The bifurcation diagram for *w^EI^* = 2.0 (i.e. after emergence of oscillations). The perturbations (±3) around fixed point for *w^EE^* = 11.5 *F*_1_ and *w^EE^* = 12.22 *F*_2_. Before bifurcation, the synaptic gating variable exhibits damped oscillations around the steady state value *F*_1_. The oscillations are sustained around the bifurcation point *F*_2_.

**Figure S6:**
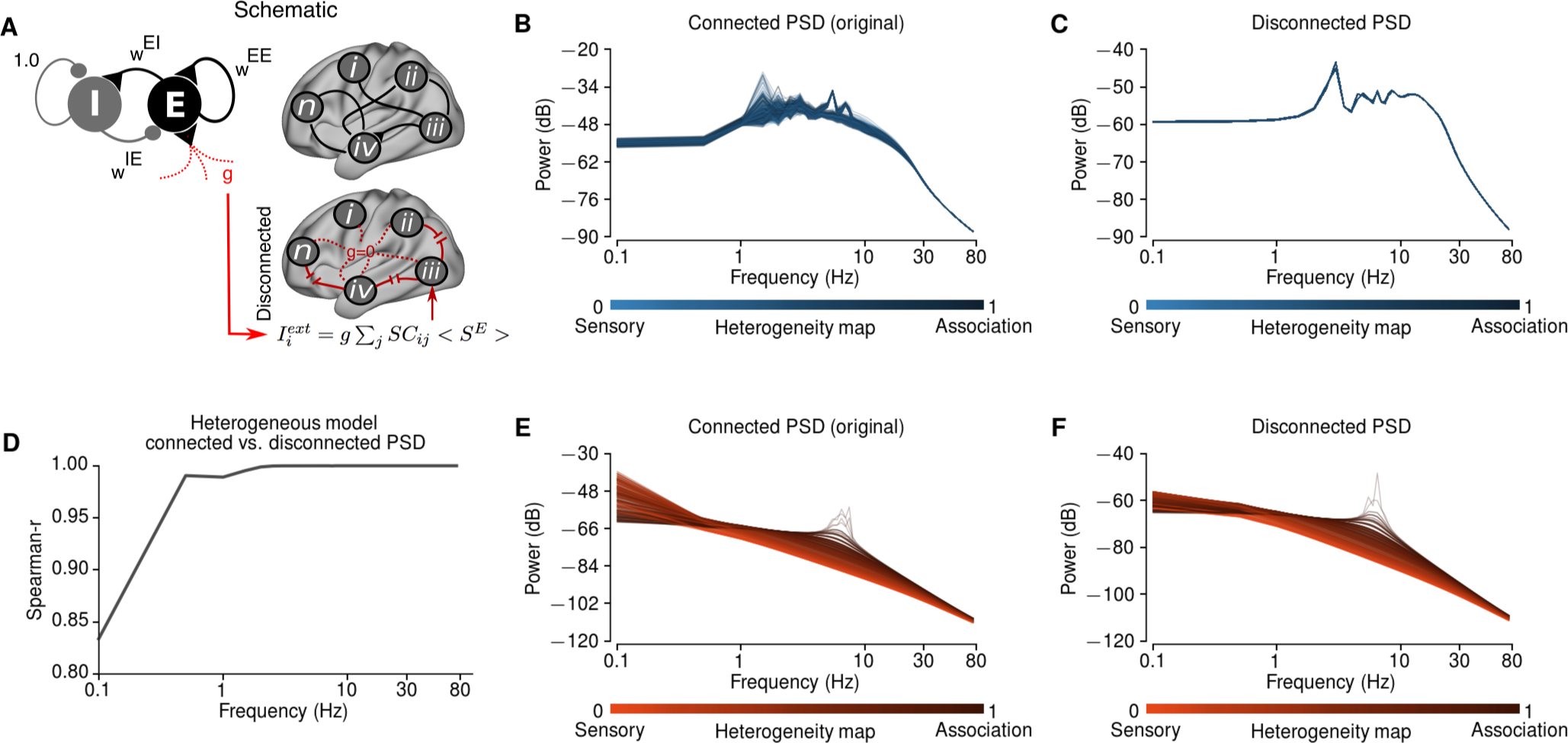
Long-Range Disconnection Analyses. **(A)** Schematic illustrating the disconnection analysis. To study the role of long-range connectivity on power spectral densities (PSDs), we calculated the power spectral density (PSD) after setting the global coupling parameter to 0 (i.e., after removing long-range connections). Since the strength of feedback inhibition (*w^IE^*) depends on the total synaptic input to each node, we added compensatory external input currents to each node such that the local microcircuit parameters were preserved. **(B–C)** PSD of the homogeneous model for the full connected model **B** and the disconnected model **C**. In the homogeneous model, the spatial patterns in the PSD were completely destroyed and collapsed into a single pattern after disconnecting the long-range connections. **(D–F)** PSD of the heterogeneous model for full connected model **E** and disconnected model **F**. Unlike the homogeneous model, the spatial patterns in the PSD were preserved in high-frequency bands (i.e. the correlation between two maps were close to 1 (**D**). The correlation between connected and disconnected PSDs were lower in very low-frequencies (i.e. <1 Hz). This shows that the regional patterns of high-frequency power emerged as a local property in the model.

**Figure S7:**
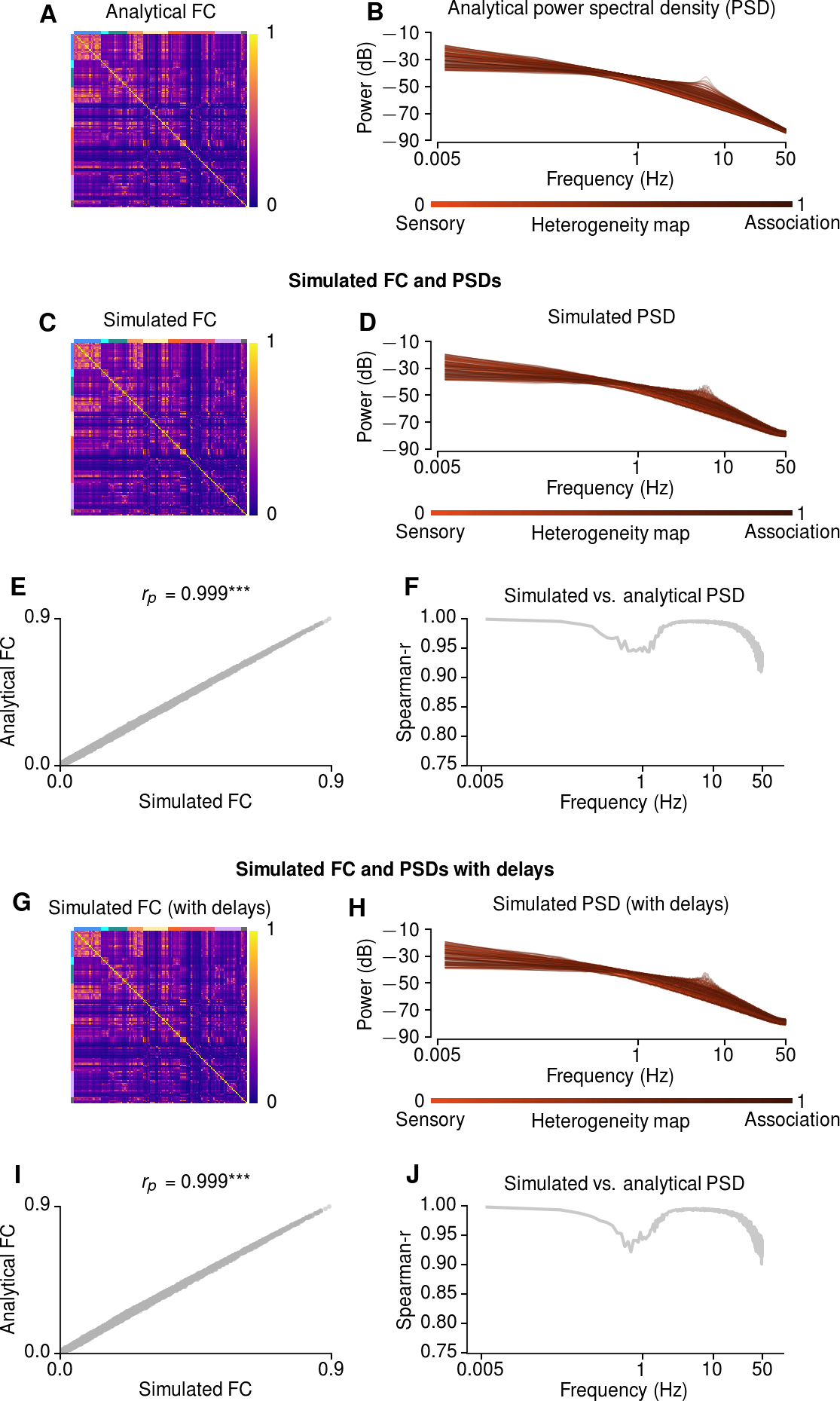
Comparison Between Numerically Simulated and Analytical BOLD FC and Power Spectral Density (PSD) in the Heterogeneous Model. **(A-B)** Analytically approximated BOLD FC **(A)** and PSD **(B)** for 20 particles drawn from posterior distribution, for the left hemisphere. FC matrices are ordered by resting state networks (marked by colored bands on top and left of matrices). **(C-F)** Numerically simulated BOLD FC **(C)** and PSD **(D)** for the same particles. The correlations between simulated and approximated values are very high for both FC (*r* = 0.999) **(E)** and PSD (*r* > 0.95) **(F)**. These results demonstrate the robustness of the analytical linearization approximation of FC. The values are averaged across 10 simulations and across all particles. *** indicates *p* < 0.0001. **(G-J)** Numerically simulated BOLD FC **(G)** and PSD **(H)** for the same particles including synaptic delays in long-range interactions due finite axonal transmission speed. The FC and PSD correlations were unaffected by long-range synaptic delays. At low frequencies related to the BOLD signal, the synaptic delays are much shorter than the characteristic timescale of the signals (see also Deco et al. (2014b)), and therefore have little impact on FC **(G)**. Synaptic delays also have little impact on the PSD patterns, at higher frequencies, in the model **(J)**. The values are averaged across 10 simulations and across all particles. *** indicates *p* < 0.0001.

**Figure S8:**
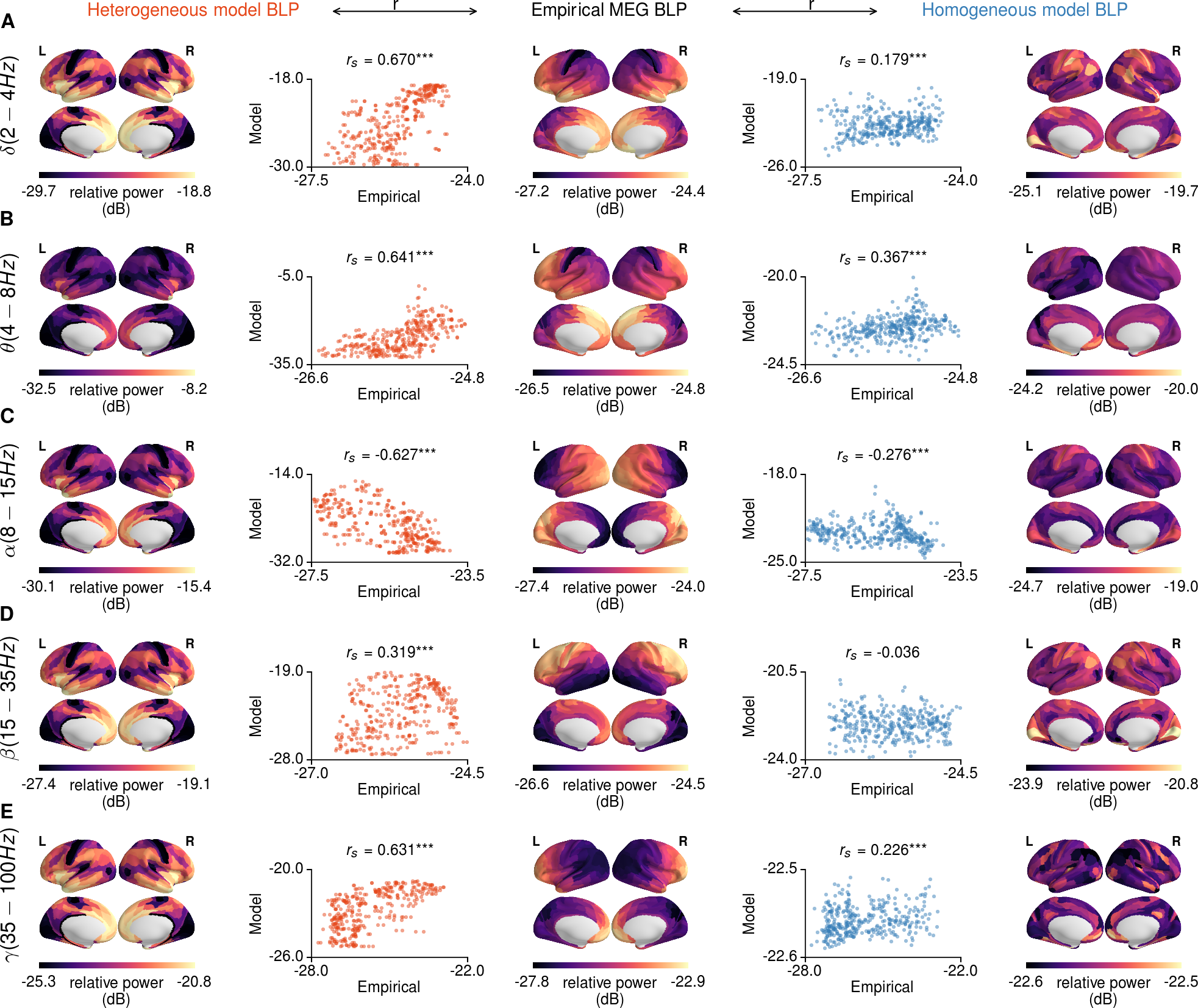
Extended Areal Maps and Correlations Between Empirical MEG BLPs and Model BLPs. **(A–E)** Spatial maps for heterogeneous model (left), empirical MEG (middle) and homogeneous model (right) BLPs. The scatter plots showing the correlations between model and empirical BLP maps are shown in between empirical and corresponding model BLP maps. The results are shown for delta (*δ*; **A**), theta (*θ*; **B**), alpha (*α*; **C**), beta (*β*; **D**), and gamma (*γ*; **E**) bands. *** indicates *p* < 0.0001 calculated via a spatial lag model.

## Acknowledgements

We thank Rishidev Chaudhuri for comments on the manuscript. This research was supported by NIH grants R01MH112746, R01MH108590, and TL1TR000141, DFG fellowship HE8166/1-1, BlackThorn Therapeutics, and the Swartz Foundation.

**Author Contributions**
Conceptualization, M.D., A.A., and J.D.M.; Methodology, M.D., J.B.B., M.H., S.N.S., A.A., and J.D.M.; Investigation, M.D., J.B.B., M.H., and J.D.M.; Resources, M.F.G., D.C.V., S.N.S., and A.A.; Data Curation, J.L.J., B.A., and A.A.; Writing – Original Draft, M.D. and J.D.M.; Writing – Review & Editing, M.D., J.B.B., M.H., M.F.G., S.N.S., D.C.V., A.A., and J.D.M.; Project Administration, J.D.M.; Funding Acquisition, A.A. and J.D.M.

